# Developing and Investigating a Nanovibration Intervention for the Prevention/Reversal of Bone Loss Following Spinal Cord Injury

**DOI:** 10.1101/2024.02.12.578222

**Authors:** Jonathan A. Williams, Paul Campsie, Richard Gibson, Olivia Johnson-Love, Anna S. Werner, Mark Sprott, Ryan Meechan, Carmen Huesa, James F.C. Windmill, Mariel Purcell, Sylvie Coupaud, Matthew J. Dalby, Peter G. Childs, John S. Riddell, Stuart Reid

## Abstract

Osteoporosis disrupts the fine-tuned balance between bone formation and resorption leading to reductions in bone quantity and quality, ultimately leading to increased fracture risk. Prevention and treatment of osteoporotic fractures is essential, for reductions in mortality, morbidity and the economic burden, particularly considering the ageing global population. Extreme bone loss that mimics time-accelerated osteoporosis develops in the paralysed limbs following complete spinal cord injury (SCI). *In vitro* nanoscale vibration (1 kHz, 30- or 90 nm amplitude) has been shown to drive differentiation of mesenchymal stem cells towards osteoblast-like phenotypes, enhancing osteogenesis, and inhibiting osteoclastogenesis, simultaneously. Here we develop and characterise a wearable device designed to deliver continuous nano-amplitude vibration to the hindlimb long bones of rats with complete SCI. We investigate whether a clinically feasible dose of nanovibration (4-hours/day, 5-days/week for 6 weeks) is effective at reversing the established SCI-induced osteoporosis. Laser interferometry and finite element analysis confirmed transmission of nanovibration into the bone, and micro-computed tomography and serum bone formation and resorption markers assessed effectiveness. The intervention did not reverse SCI-induced osteoporosis. However, serum analysis indicated an elevated concentration of the bone formation marker procollagen type 1 N-terminal propeptide (P1NP) in rats receiving 40 nm amplitude nanovibration, suggesting increased synthesis of type 1 collagen, the major organic component of bone. Therefore, enhanced doses of nanovibrational stimulus may yet prove beneficial in attenuating/reversing osteoporosis, particularly in less severe forms of osteoporosis.

---

Osteoporosis is a worldwide public health concern of increasing importance due to an ageing population.^1^ It is a progressive metabolic bone disease, that reflects a disruption in the fine-tuned balance between the coupled processes of bone resorption and formation, favouring resorption. Bone quantity and quality progressively depreciate, elevating susceptibility to fracture which is associated with increased morbidity, mortality and healthcare costs.^2^ In the UK, the economic burden of osteoporotic fracture is £4 billion per year, while in the US this is $17.9 billion.^3^ The prevention and treatment of these fractures is of paramount importance to society.

Strategies currently available for attenuating or preventing the development of osteoporosis address the imbalance on one-side, that is, attenuating bone resorption (*i.e.* bisphosphonates, receptor activator of nuclear factor-κB ligand (RANKL) antibody and selective oestrogen receptor modulators (SERMs)) or enhancing bone formation (*i.e.* teriparatide and abaloparatide).^4^ This strategy is suboptimal, as existing microstructural deterioration and hence fragility is not reversed in anti-resorptive strategies, while anabolic strategies can partly restore microstructural deterioration, and there is some evidence that anabolic agents may reduce fracture risk more effectively.^5^ Another strategy is the dual-action approach, a combination of anabolic and antiresorptive strategies either successively or together which could reduce fracture risk more than either strategy alone.^5^

*In-*vitro nanoscale vibration (1 kHz, 30 – 90 nm amplitude) applied continuously for up to 4-weeks has been demonstrated to differentiate adult human bone marrow-derived mesenchymal stem cells (MSCs) towards the osteoblast cell linage, in both 2D culture and 3D soft-gel constructs, without aid from osteogenic scaffolds or chemicals.^6–8^ Further, osteogenesis has been confirmed by the occurrence of mineralisation within soft-gel constructs.^8^ Furthermore, nanovibrational stimulation of 3D co-cultures of primary human osteoprogenitor and osteoclast progenitor cells simultaneously inhibits osteoclastogenesis and enhances osteogenesis.^9^ This nanoscale vibration is supplied by a bespoke nano-amplitude vibrational bioreactor.^10^

To translate this technology for direct *in-vivo* applications, the vibration delivery platform needs to be miniaturised into a non-invasive, wearable configuration. Secondly, a suitable animal model is needed to test its efficacy. Osteoporosis is commonly associated with ageing and the post-menopause. However, an extreme form of osteoporosis is observed at the ends of the long bones (around the knee) following complete spinal cord injury.^11^ Rat models of complete spinal cord injury (SCI) are time accelerated models of bone loss which replicate the bone loss observed in the human SCI population.^12^

There are other vibration interventions that have applications in bone health. The two main interventions being whole body vibration (WBV)^13^ and low intensity pulsed ultrasound (LIPUS),^14^ each has some similarities with nanovibration that they do not have with each other. For WBV, the typical peak acceleration is between 0.3 and 0.6 g, which is comparable to the peak accelerations produced by the nanovibration used here (0.2 to 0.4 g), however, the vibration parameters are significantly different (typically <100 Hz, >1 mm). Another similarity is that the vibration is delivered continuously (not pulsed) and sinusoidally. However, the delivery mechanism of vibration to the bone (and bone cells) is decidedly different. In WBV, the stimulus is designed to be delivered to the bone through vibration-induced muscle driven dynamic stimulation^13^. The higher frequency of nanovibration would suggest that muscle fibres are unresponsive^15^. LIPUS on the other hand, is a targeted pulsed oscillation that penetrates into the bone tissue. Reports indicate it has a beneficial role in fracture healing,^14,16^ and recent experimental evidence indicates that it can promote MSC differentiation towards osteoblast lineages and mineralisation.^17,18^ The most effective parameters of LIPUS for fracture repair are at a pulse excitation frequency of 1.5 MHz, intensity (spatial average temporal average) of 30 mW/cm^3^, duty cycle of 20% and pulse repetition frequency (PRF) of 1 kHz.^14^ We note the correspondence between the PRF and the frequency of nanovibration used here. Both WBV and LIPUS have demonstrated some potential for the attenuation of mild osteoporosis in animal models.^19–22^ However, this does not include SCI induced osteoporosis, where the effects of WBV remain unclear,^23,24^ while the application of LIPUS to the calcaneal bone for 6 weeks in people with SCI showed no effect.^25^

The overall aim of this study was to investigate the efficacy of nanovibration (1 kHz frequency, 40 or 100 nm amplitude) at reversing (or preventing) the SCI-induced osteoporosis observed in the paralysed hindlimbs of completely spinal cord transected rats. The specific objectives were to; i) develop, test and characterise the effectiveness of a device that delivers nanovibration to the hindlimb long bones of spinal cord transected rats, and; ii) determine the effect(s) that unilateral nanovibration of two different amplitudes - 40 nm and 100 nm - has on the induced osteoporosis.

## Results & Discussion

### Development of a wearable nanovibration delivery device

*In-vitro* nanovibration-induced bone mineralisation is observed in human bone marrow-derived MSCs after 4 to 6-weeks of continuous exposure (24 hours/day).^8^ It is not feasible to continuously vibrated rat hindlimbs. To confirm that osteogenesis can occur with non-continuous (<24 hour/day) nanovibration, and to determine a feasible vibration dose for *in vivo* applications, MG63s (an osteoblast-like cell line) and MSCs (from human bone marrow) were stimulated at 1 kHz with 30 nm or 90 nm amplitude vibration either continuously or for 4 hours per day (intermittent group), and compared static control. After 14-days of MG63 culture, quantitative real time PCR (qRT-PCR) revealed that the expression of the early stage osteogenic marker runt-related transcription factor 2 (RUNX2) was significantly upregulated in 30 nm and 90 nm amplitude intermittently vibrated groups, and 30 nm amplitude continuously nanovibrated group compared to static control, indicating that a key transcription factor associated with osteoblast differentiation is upregulated (Figure 1A). Further, no significant differences were observed in osteogenic gene expression between continuous or intermittent nanovibration at either time point (Figure 1A). Furthermore, after 28-days of MSC culture, no significant differences were observed in early (RUNX2 or collagen I (COL1A)) or later stage osteogenic markers (osteonectin (ON), alkaline phosphatase (ALP) or osteocalcin (OCN)) between continuous vibrated, intermittently vibrated MSCs or non-vibrated controls (Figure 1B). This agrees with our previous work, that showed no differences in these markers after 14 days of continuous vibration, indicating that at this stage the osteogenic process is transcriptionally complete.^8^ This data suggests that an intermittent stimulation regime may be suitable to provide a comparable osteogenic stimulus *in vivo*.

**Figure 1.**
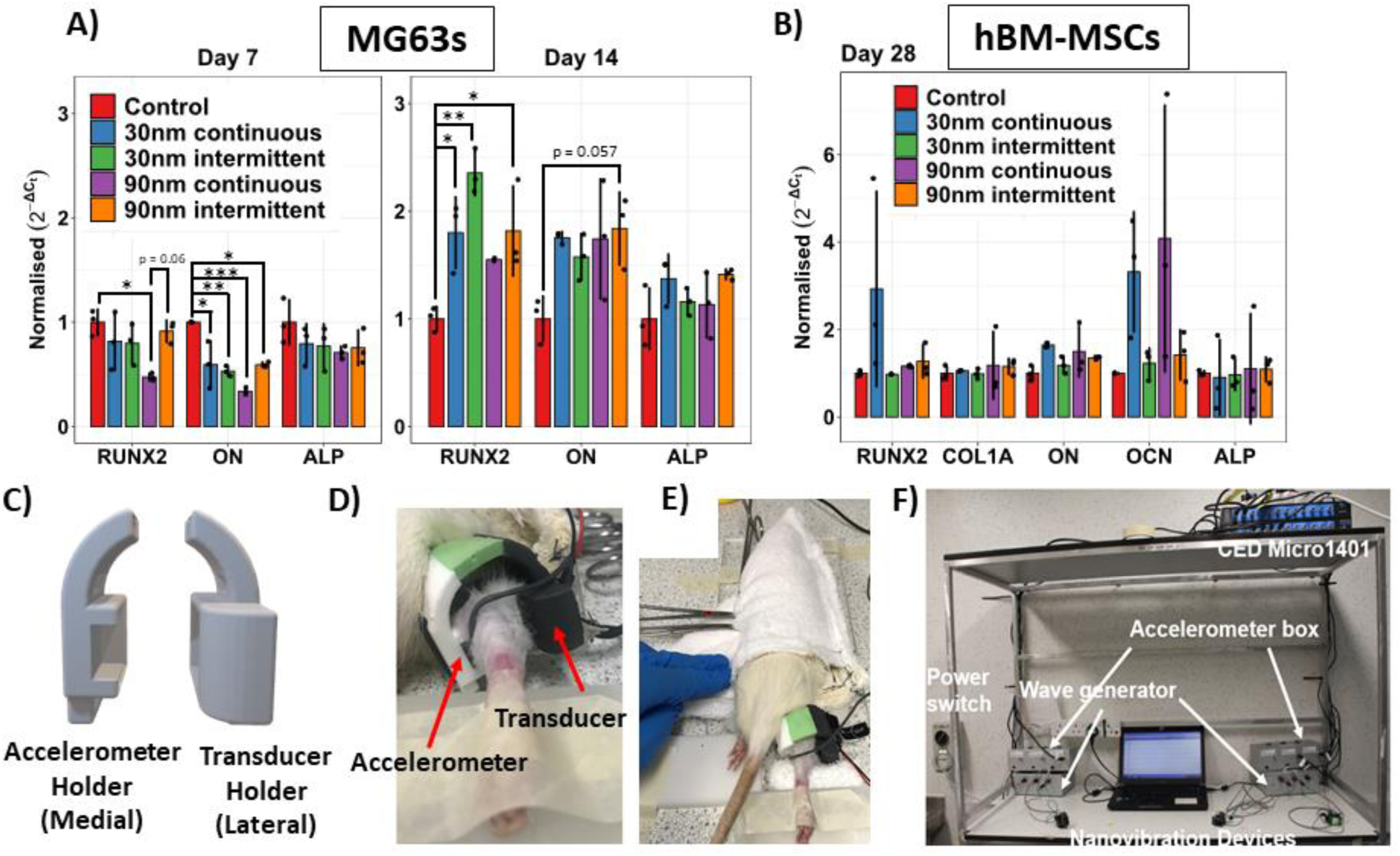
Development of a wearable nanovibration delivery device. A) 7- and 14-day qRT-PCR analysis of RUNX2, ON and ALP transcripts in MG63 cells comparing continuous and intermittent (4-hours per day) doses of nanovibration at 30 nm and 90 nm. B) 28-day qRT-PCR analysis of RUNX2, COL1A, ON, OCN and ALP in human bone marrow-derived MSCs comparing continuous and non-continuous (4-hours per day) doses of nanovibration at 30 nm and 90 nm. C) CAD drawing of the 3D-printed housing of the transducer and accelerometer. D) A close-up of the nanovibration delivery device attached around the knee of the right hindlimb of a spinal cord transected rat. E) The rat is lightly restrained within a soft towel pouch for the duration of each intervention session. E) Overview of the complete experimental setup.

With a valid nanovibration duration of 4-hours per day, the nanovibrational bioreactor technology^10^ was translated into a wearable form. Specifically, devices were developed to deliver nanoscale vibration to the MSCs within the bone marrow of the hindlimbs of awake spinal cord transected rats. The nanovibration delivery device and associated electronic systems were designed, manufactured and validated in-house specifically for this study (Figure 1C – F). The device consisted of a bone conduction transducer and accelerometer housed within a custom-made 3D-printed plastic harness, that were strapped directly to the hindlimb at the lower knee (Figure 1C - E). For more information regarding the design of the device see Supplemental Information 1. The devices were designed to meet multiple experimental needs, including: i) to deliver continuous sinusoidal nanoscale mechanical oscillation at a frequency of 1 kHz just below the knee to the trabecular-rich proximal tibia in the paralysed hindlimbs of spinal cord transected rats, ii) to measure and log the transmitted vibration, iii) to provide the operator in real-time the amplitude of the transmitted vibration, thus enabling adjustment of the amplitude in real-time, to ensure it remained within acceptable pre-defined limits. The design, fit and optimisation were refined through multiple iterations on cadavers. The advantage of using the rat model of complete spinal cord transection as the model of bone loss is that the resulting permanent and complete hindlimb paralysis and concomitant loss of sensation, meant that these rats tolerate well the direct application of a device directly to the hindlimb for a prolonged period of time. If another model of bone loss was used then the animal would need to be anaesthetised for the duration of each 2-hour intervention.^26^ During the intervention, the rat was lightly restrained with a soft-towel pouch (Figure 1E). The device was attached unilaterally to the right hindlimb only (Figure 1D and E). The attachment of the device required that the hindlimb be taped down with the tibio-femoral angle at approximately 120 degrees. Our setup enabled multiple rats to go through the intervention simultaneously. Calibration of the devices was carried out weekly during the intervention (Supplemental Information 2).

### Characterisation of the transmission of vibration to hindlimb long bones

Prior to commencing the nanovibration intervention, the transmitted vibration amplitude at 1 kHz was measured in anesthetised rats, using laser interferometry, to confirm that suitable nanovibration parameters are deliverable to the hindlimb long bones. To optimise the delivery of nanovibration, this needed to be tested in the presence of muscle wasting, so spinal cord transected rats were used (*n* = 3). Transmitted vibration amplitudes were measured on the anteromedial surface of the right proximal tibia, which was surgically exposed under general anesthesia. For these measurements, the device’s accelerometer was also attached on the skin’s surface just below the exposed bone (Figure 2A). That is, two independent measures of the transmitted 1 kHz vibration amplitude were acquired simultaneously, one on the bone’s surface and one on the skin’s surface.

**Figure 2.**
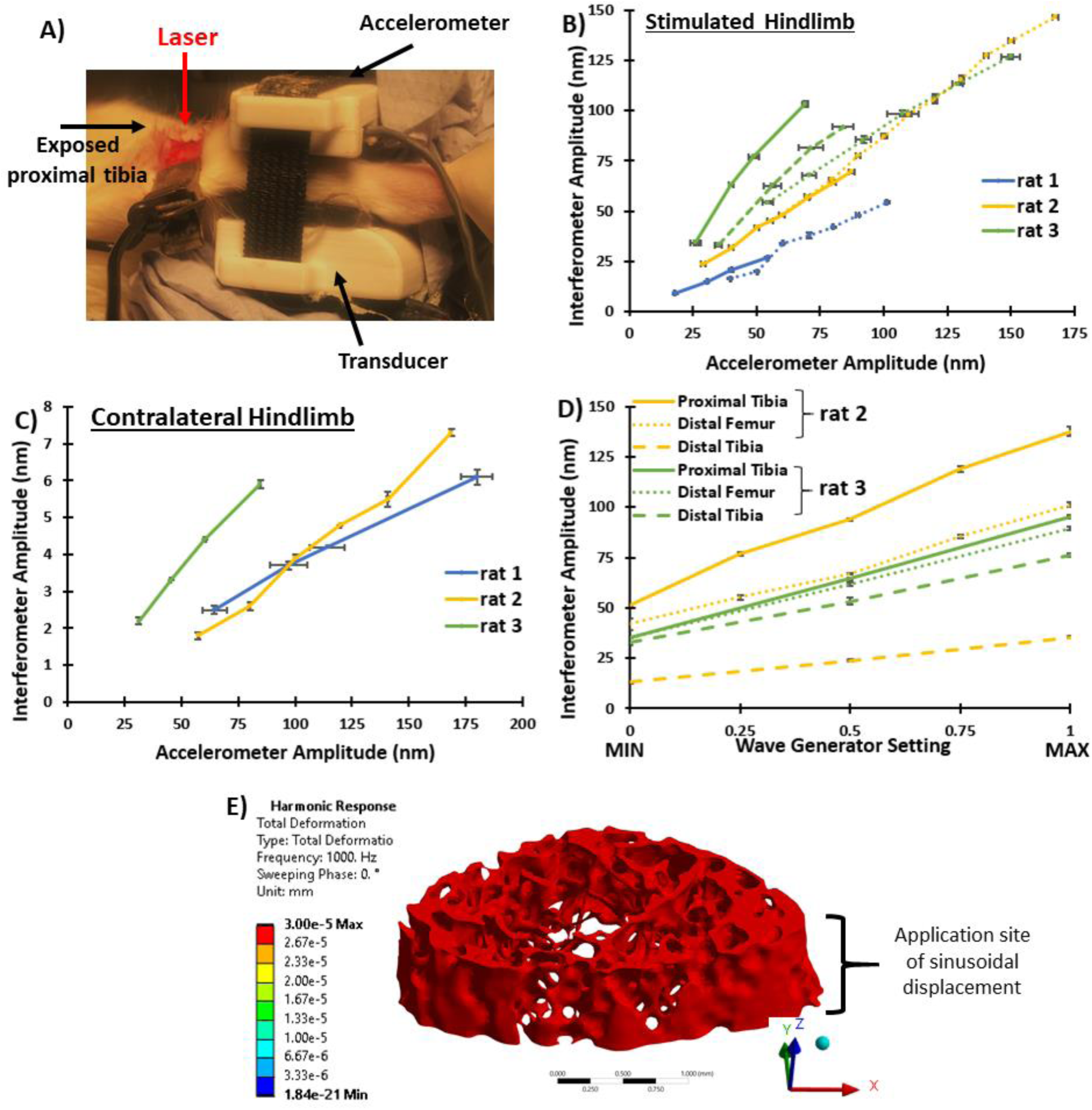
Transmission of nanovibration through bone. A) Exposed stimulated proximal tibial laser interferometric measurement site in relation to transducer and accelerometer. B) Plot of interferometer-derived transmitted vibration amplitude from the stimulated proximal tibial bone surface against accelerometer-derived transmitted vibration amplitude from the skin’s surface, for three spinal cord transected rats. Patterned lines indicate separate measurements where the device was removed and reattached. C) Plot of interferometer-derived (from exposed contralateral proximal tibia) and accelerometer-derived amplitude (from vibrated proximal tibia) for the different rats. D) Plot of interferometer-derived amplitude from multiple exposed bone sites on the vibrated hindlimb. E) Harmonic response finite element analysis of the distal femoral metaphyseal trabecular bone, showing the predicted rigid-body like transmission of nanovibration (minimal internal deformation). 3.00e-5 mm = 30 nm. Data shown as mean ± SD.

Accelerometer-derived vibration amplitude measurements of the stimulated hindlimb were overall proportional to those measured with laser interferometry at the proximal tibial bone surface (Figure 2B). However, a variation exists between the accelerometer- and interferometer-derived amplitude, for example a 40 nm accelerometer-derived amplitude measurement translated to between 18 nm and 62 nm amplitude at the vibrated proximal tibial bone surface (Figure 2B). This is partially explainable by differences in rat hindlimb musculature between individual rats and also variability in attachment of the device. This variability is depicted for each rat with the sets of patterned line, where between each of these measurements the device was completed removed and then reattached (Figure 2B). These results indicate that the device must be attached to the hindlimb in a reproducible manner. The spread of nanovibration to the contralateral (i.e. non-stimulated) hindlimb was also measured at the exposed proximal tibia, importantly it was found to be minimal (Figure 2C). All vibration amplitudes measured at the non-stimulated hindlimb were below the *in vitro* determined osteogenic threshold of nanovibration (20 – 100 nm) ^7^, even when the stimulated hindlimb experienced vibration amplitudes several times higher than the lower limit of this threshold (Figure 2C). This provides confidence that effective nanovibration was delivered to the directly stimulated, but not to the contralateral hindlimb long bones.

Further, the propagation of nanovibration along the length of the stimulated hindlimb was measured at multiple exposed bone sites with laser interferometry only (Figure 2D). This indicated that nanovibration was transmitted throughout the length of the tibia and femur, but diminished with distance from the transducer. Throughout the stimulated hindlimb long bones, vibration amplitudes at 1 kHz were observed that could be considered osteogenic (Figure 2D).

Finally, finite element analysis (FEA) software (ANSYS 2022 R2) was used to simulate and predict how nanovibration propagates through the trabecular bone of the distal femur, by performing a harmonic response analysis with the bone structure based on µCT data. The vibration (1 kHz, 30nm) was applied from the lateral (negative *x*) direction over a surface area that replicates the positioning and size of the vibration transducer. The simulation shows that the vibration propagates through the trabecular bone in a rigid body-like manner, that is it propagates with no internal stresses, causing no deformation (Figure 2E). This suggests that the vibration is fully transferred through the width of the bone with minimal attenuation.

The transmission of nanovibration to bone is then significantly different to both WBV and LIPUS. In WBV, the stimulus is delivered to the bone through induced muscle-driven dynamic stimulation, while the higher frequency of nanovibration would suggest that muscle fibres are unresponsive.^15^ While both nanovibration and LIPUS can be described as an alternating pressure wave, the higher frequency components of LIPUS indicate that it generates fluctuating pressure within tissue,^27^ thus reflection and attenuation significantly affect transmission. In fact, when ultrasound propagates into intact bone up to 40% of the energy is reflected at the soft tissue-bone boundary.^28^ Furthermore, greater than 80% of the remaining energy is attenuated within the first millimetre of cortex.^28^ Therefore, it can only be guaranteed that periosteal cells on the surface of the cortex in the region of the LIPUS application are biophysically stimulated. Interestingly, dissected rat femora exposed to LIPUS exhibited a site-specific periosteal effect.^29^ Specifically, at the angle of LIPUS application only, increased periosteal mineralisation was observed.^29^ The laser interferometric and FEA analyses performed here suggest that the much larger wavelength of nanovibration (approximately 3.5 m for cortical bone) compared to the dimensions of the bone (< 1 cm diameter of metaphysis) results in a very minimal pressure change over the bone, suggesting that the vibrational behaviour of the hindlimb long bones at 1 kHz is likely rigid-body motion, with the directly stimulated tibial bone moving pistonically (in unison). Assuming vibration in a rigid body manner this suggests that the encapsulated trabecular bone was also being nanovibrated within the osteogenic *in vivo* range, as verified by FEA (Figure 2E). Interestingly, LIPUS (pulse excitation frequency of 1.5 MHz, intensity (spatial average temporal average) of 30 mW/cm^3^, duty cycle of 20%, PRF of 1 kHz) applied to exposed bone from a fractured cadaveric human forearm has been shown to induce 1 kHz motion at the nanoscale.^30^ A hypothesis has previously been made that this low frequency (1 kHz) radiation force, not the higher frequency (1.5 MHz) pulsed ultrasound, is responsible for at least some of the observed biological effects.^31,32^ *In vitro* experiments of ATDC5 chondrocytes showed that treatment with a 1 kHz square wave at 20% duty cycle induced chondrogenesis similar to treatment with 1.5 MHz LIPUS.^31^ Furthermore, varying the PRF (1-, 100-, 1000 Hz) of LIPUS led to differential responses in the calcium secretion of bone marrow derived MSCs, with increased response at the higher frequencies.^33^ Cells appear sensitive to the PRF and the influences that the ultrasonic and acoustic radiation (PRF) force components of LIPUS have on the osteogenic response still remains an open question.^14^

### Investigating nanovibration to reverse established spinal cord transection-induced osteoporosis

The rat model of complete spinal cord transection-induced osteoporosis^12,26^ was used to investigate the efficacy of nanovibration at reversing established induced osteoporosis. Two amplitudes, 30 nm and 90 nm, have both been shown to induce osteogenesis *in vitro*, with the higher amplitude producing the greater osteogenic response.^7^ Based on the limitations of the driving electronics, two amplitudes within a similar range to the *in vitro* studies were investigated, 40 nm (N40) and 100 nm (N100), as measured by the device’s accelerometer. 6-weeks were allowed to pass from the time of spinal cord transection surgery to the start of the nanovibration intervention, to allow time for significant trabecular bone loss, replicating that seen in chronic SCI-induced osteoporosis.^12,26^ Nanovibration was then applied continuously for two 2-hour sessions/day, 5-days/week for 6-weeks. The intervention lasted 6-weeks to coincide with the average bone turnover period in rats, which is approximately 40 days.^34^ Age-matched spinal cord transection (SCI) and sham-operated control (AGE-CTR) rats were also used for comparison. Initiating the nanovibration intervention at an earlier time post-surgery (3-days) was also investigated in a preliminary study to test the efficacy of nanovibration at attenuating ongoing SCI-induced osteoporosis (Supplementary Information 3).

There was no difference in body mass between groups at time of surgery (Supplementary Information 4). From day 3 post-surgery and onwards AGE-CTR body mass was higher than all other groups (*p* < 0.05) (Supplementary Information 5). There was no difference in the body mass between N40, N100 and SCI groups at any time point post-surgery. There was no difference in gastrocnemius muscle mass at the end of the intervention between left and right hindlimbs for any group (Supplementary Information 6). Also, no differences were detected in gastrocnemius mass between N40, N100 and SCI groups suggesting that nanovibration does not stimulate muscle fibres.

Trabecular bone was evaluated by micro-computed tomography (µCT) (Figure 3). In the proximal tibial metaphyseal trabecular bone, the region directly nanovibrated, there were no significant improvements in bone quantity or microarchitecture in either N40 or N100 vibrated hindlimbs when compared to contralateral control (Supplementary Information 7) or when compared to each other or with SCI rats (Figure 3A). Overall, a similar scenario is described for the proximal tibial epiphyseal and the distal femoral metaphyseal and epiphyseal trabecular bone (Supplemental Information 8). This was also the case for nanovibration intervention started 3-days post-surgery (Supplemental Information 3). Additionally, there was no change in the orientation of trabecular bone as measured by the degree of anisotropy (DA) in the proximal tibial metaphysis trabecular bone (Figure 3B). The observation that nanovibration propagated throughout the vibrated hindlimb long bones (Figure 2D), not just in the regions directly stimulated by the transducer, motivated a global survey of the trabecular bone. However, no differences in trabecular BA/TA were observed at any point along the tibia’s length, data shown here for vibrated and contralateral control tibia of N40 rats only (Figure 3C). Cortical bone was also evaluated with µCT and mechanical testing with three-point bending. Nanovibration had no effect on cortical bone (Supplemental Information 9 and 10).

**Figure 3.**
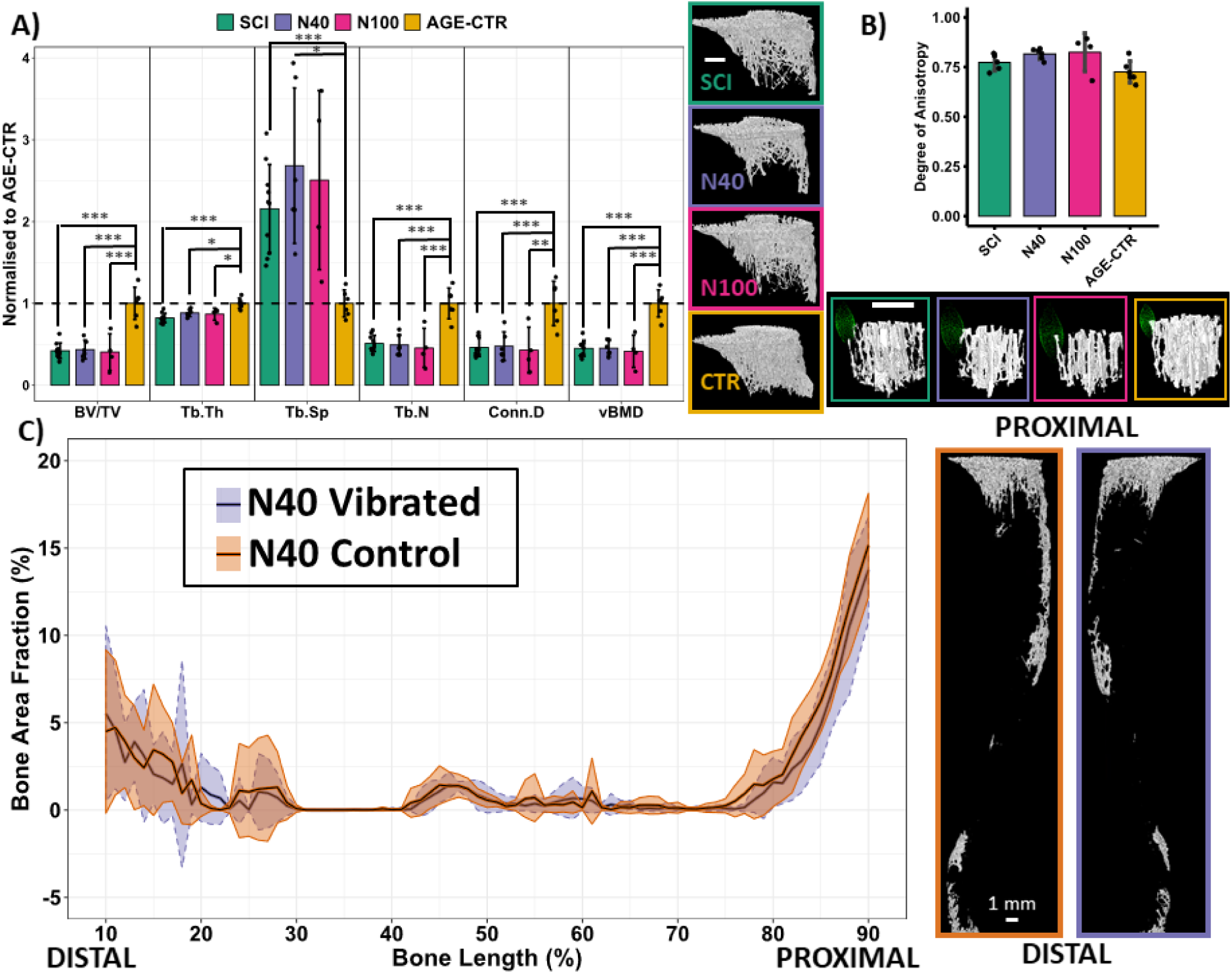
Effect of nanovibration intervention on trabecular bone quantity and microarchitecture. A) µCT-derived morphometric and densitometric analysis and representative images of proximal tibial metaphyseal trabecular bone; the region directly stimulated by the device. The parameters measured being bone volume fraction (BV/TV), trabecular thickness (Tb.Th), trabecular separation (Tb.Sp), trabecular number (Tb.N), connectivity density (Conn.D) and volumetric bone mineral density (vBMD). B) µCT-derived degree of anisotropy analysis of a trabecular bone cube within the proximal tibial metaphysis. Representative bone cube and point cloud of mean intercept lengths shown. C) Global survey of trabecular bone area fraction for N40 rat vibrated and contralateral whole tibia (excluding epiphyses), and representative µCT-based images. White scale bars indicate 1mm. Data shown as mean ± SD.

Serum bone formation and resorption were measured using procollagen type 1 N-terminal propeptide (P1NP) and C-terminal telopeptide of type I collagen (CTX), respectively, immediately following the end of the intervention (Figure 4A). The concentration of the gold standard bone formation serum marker P1NP was found to be elevated by 67% (*p* < 0.01) in the N40 group relative to the SCI group at the end of the intervention. No differences between groups were observed for the bone resorption marker CTX. These results suggest that nanovibration of certain amplitudes increases bone formation processes (synthesis of type 1 collagen) without negatively affecting bone resorption.

**Figure 4.**
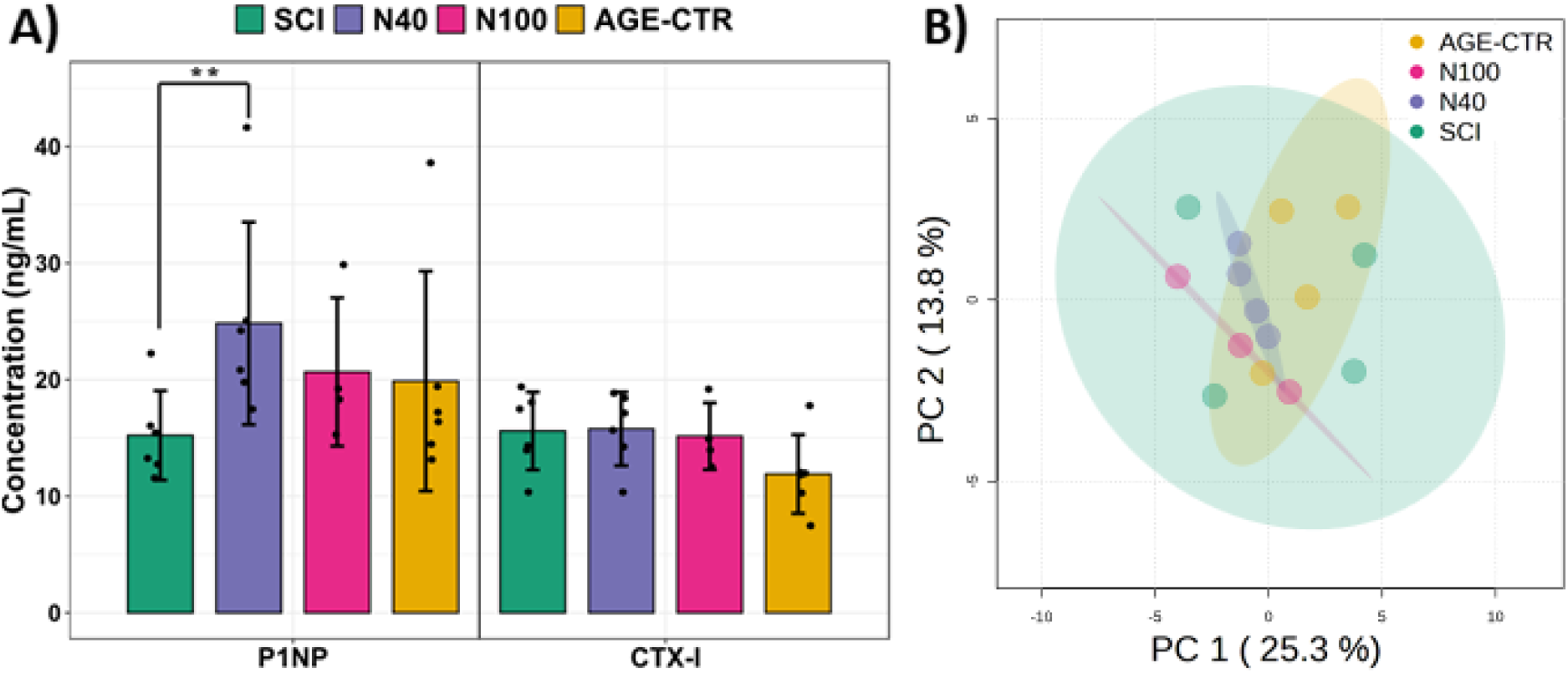
A) Serum levels of bone turnover markers in SCI, N40, N100 and AGE-CTR groups for bone formation marker procollagen type 1 N-terminal propeptide (P1NP) and bone resorption marker C-terminal telopeptide of type I collagen (CTX). Data shown as mean ± SD. ** indicates *p* < 0.01. B) Principal components analysis (PCA) of blood serum-derived metabolites.

To investigate further the cellular response of the targeted nanovibration intervention, we took an untargeted metabolomics approach. Blood serum-derived metabolites were analysed by hydrophilic interaction liquid chromatography-mass spectrometry. Principal component analysis (PCA) revealed system level changes (Figure 4B). Despite no separation of the first two principal components, the whole metabolome was differentially regulated. SCI caused an increase in metabolite variance, particularly in the first principal component (PC 1). However, nanovibration reduced this variance, most noticeably for the N40 group. Further metabolomic pathway analysis did not reveal information regarding bone turnover.

### Nanovibration delivered to bone during intervention

The inclusion of a calibrated accelerometer is a key feature of the device. It was included to allow the operator to monitor and control the transmitted vibration parameters in real time, so that vibration was continuously delivered in a consistent manner. The recorded vibration amplitude at 1 kHz from a representative 2-hour intervention session, and the average peak transmitted amplitude for all such sessions for a representative N40 rat are shown in Figure 5A and B, respectively. This shows that during each session rats received a relatively consistent amplitude of vibration. Only one rat received the maximum intended duration of the intervention (Figure 5C). The remaining nanovibrated rats received a range of durations from 33% up to 97% of the intended dose. Overall, there was a larger variation in both the amplitude of transmitted nanovibration and total vibration time between N100 rats than between N40 rats (Figure 5D). The added level of precision and reproducibility that measuring the transmitted vibration provides is most often missing in studies that assess vibration’s ability at increasing bone mass or density. For whole body vibration (WBV), it is recommended that the platform’s vibration parameters are measured with an accelerometer prior to the start of an intervention,^35^ however, this is an extremely rare occurrence. Attenuation (or amplification) of this waveform however means that it is highly probable that the vibration parameters transmitted to the bone regions under investigation are significantly different to the vibration parameters measured on the platform.^36^ In our study we observed that slight variations in the contact of the device with the rat hindlimb can lead to significant changes in the vibration dose transmitted. To our knowledge, the measurement of transmitted vibration to bone in LIPUS studies has yet to be attempted.

**Figure 5.**
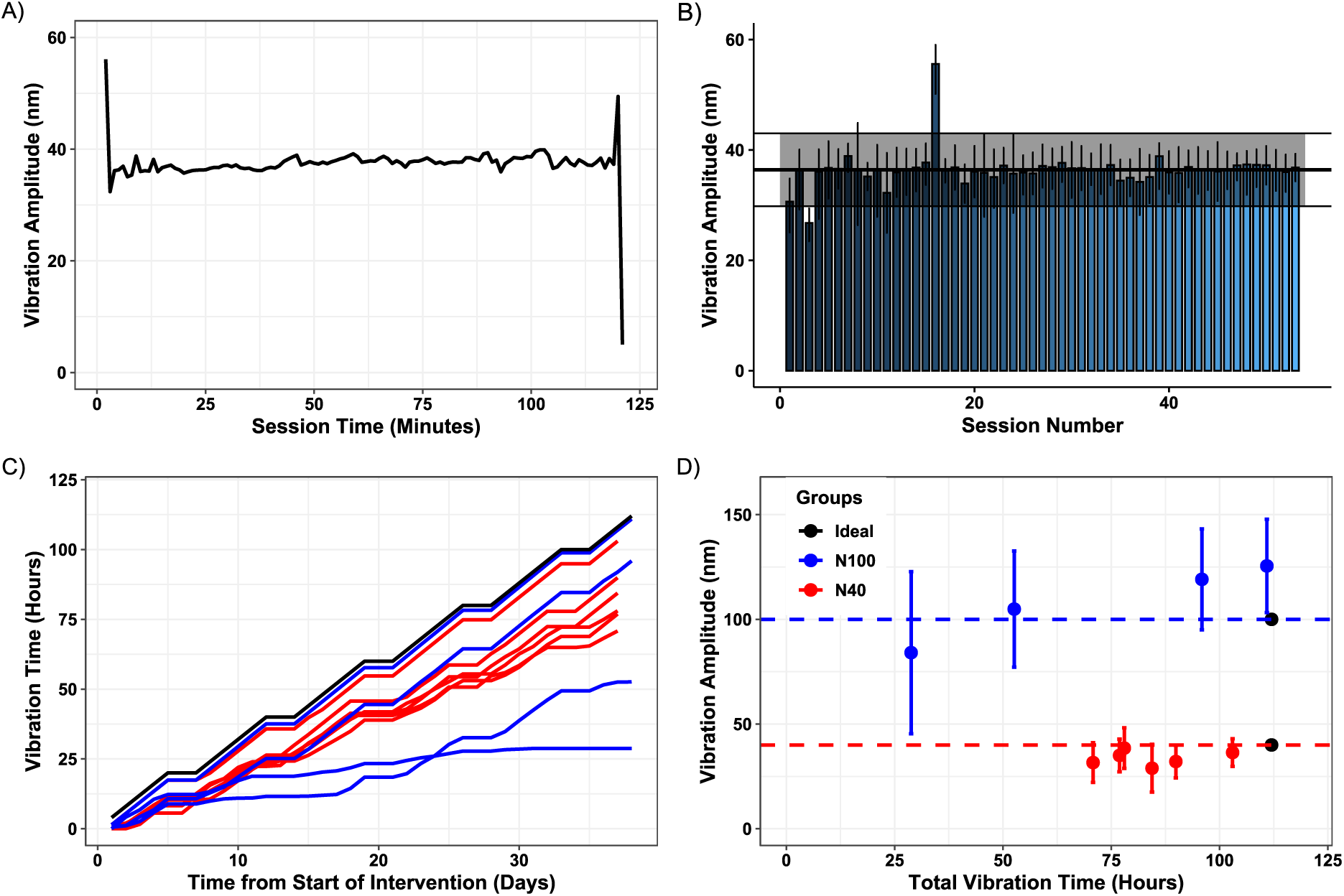
Transmitted nanovibration data summary. A) Accelerometer-derived amplitude data from a representative rat. B) Average amplitude ± SD per nanovibration intervention session for this rat. Total average displacement ± SD throughout entire intervention is superimposed on top. C) Cumulative plot of vibration time for each rat per day from start of intervention. D) Plot of vibration amplitude ± SD versus overall vibration time, summarising the nanovibration exposure for each rat, with the ideal combination of both for N40 and N100 groups plotted as black dots.

Despite not reversing established SCI-induced osteoporosis, this study provides the first evidence in support of the use of a device and experimental setup that can deliver a vibrational stimulus targeted specifically at the hindlimbs of a paralysed rodent model for prolonged periods of time (>20 minutes) without the use of anaesthesia. This has potential use for testing a variety of vibration parameters as well as other types of biophysical stimulation as therapies for bone loss.

There are several potential hypotheses to explain why this specific nanovibration intervention did not reverse existing SCI-induced bone loss. Firstly, the vibration stimulus intensity, which is a function of duration, amplitude and frequency, may not have been sufficient to replace SCI-induced bone loss. Further work is needed to determine if other nanovibration dose parameters provide an osteogenic effect *in vivo*. Secondly, human studies have noted that the skeletal system in patients with chronic SCI appears to be resistant to change in response to electrical muscle stimulation and WBV.^37,38^ These studies concluded that preventing SCI-induced osteoporosis may be more effective than reversing the established (chronic) condition. Another consideration is whether applying the vibration for a longer overall duration (several bone remodelling cycles) would be more effective. It is also possible that an alternative model, such as, ovariectomy-induced osteoporosis (OVX), ^39^ where the bone loss is milder and less rapid, would reveal effects not seen in the more severe SCI model. However, the main disadvantage of a model without hindlimb paralysis (*e.g.* OVX) is that it is challenging to design a device that awake animals will tolerant wearing for long enough periods of time that would deliver a sufficient dose. For these small animal models, an untargeted vibration delivery mechanism (*e.g.* a whole cage approach) may be required.

Our study had several limitations. Rats did not tolerate attachment of an inactive device to the contralateral control hindlimb. We also did not set up a control spinal cord transected rat group that received unilateral attachment of the device without stimulation applied. Either of these measures would have controlled for any unintended effects related to the attachment of the device itself. The second option would also have ensured that the unilateral application of nanovibration did not produce any systemic effects. However, the quantification of the transmission of nanovibration to the contralateral hindlimb (Figure 2C) provided confidence that this control hindlimb was not receiving the nanovibration stimulus. A further improvement would have been to perform dynamic histomorphometry and tartrate-resistant acid phosphatase staining to obtain information regarding bone formation and bone resorption, respectively. This would provide more sensitive insights into the cellular response to nanovibration.

## Conclusions

In this study, a clinically feasible dose of intermittent nanovibration (4 hours per day) was identified that produces equivalent osteogenic effects in an osteoblast-like cell line and human bone marrow-derived MSCs to that of continuous nanovibration (Figures 1A and B). This meant that a wearable nanovibration delivery device and intervention could be developed (Figures 1C – 1F). Laser interferometry and FEA were utilised to demonstrate that suitable nanovibration (1 kHz, 30 nm – 90 nm) was deliverable to the trabecular bone within the proximal tibia (Figure 2). This was followed by an investigation of a nanovibration intervention for the reversal and/or prevention of bone loss following complete spinal cord transection in rats, which produces a very severe, but reproducible bone loss within the paralysed hindlimbs. Nanoscale vibration, that inhibits osteoclastogenesis and enhances osteogenesis *in vitro,* were delivered to the paralysed hindlimb long bones in a continuous and consistent manner (Figure 5). This protocol did not reverse or attenuate the induced osteoporosis (Figure 3). However, blood serum analysis indicated an elevated concentration of the bone formation marker P1NP in rats receiving the 40 nm amplitude intervention (Figure 4). This suggests that nanovibration increased the synthesis of the main component of the organic matrix of bone - type 1 collagen. Other doses of nanovibration stimulus may yet prove productive at attenuating or reversing bone loss, particularly in less severe types of osteoporosis.

## Methods

### Cell Culture

Human bone marrow mesenchymal stem cells (MSCs, PromoCell) and MG63 cells (ECACC) were cultured separately in Dulbecco’s modified essential medium (DMEM, Sigma), supplemented with 10% (v/v) fetal bovine serum (FBS, Sigma), 1% non-essential amino acid (MEM NEA, Gibco) and 2% antibiotics (penicillin/streptomycin, Sigma). MSCs were used at passage 4 and MG63s at passage < 20. MG63 cells were seeded at 1136 cells/cm^2^ and MSCs by 4000 cells/cm^2^ into 96-well plates. MG63s were seeded at a lower density to avoid overgrowth by day 14. Cells were incubated at 37°C with 5% CO2 and media changed every 3 days.

96-well plates were magnetically-coupled to the bespoke nano-amplitude vibrational bioreactor to receive nanovibration stimulation. Nanovibration stimulation started 24 hours following seeding to allow cells to adhere. Control cells were cultured without nanovibration stimulation. Nanovibrated cells were either continuously nanovibrated at an amplitude of 30 or 90 nm, or intermittantly vibrated at 30 or 90 nm for 4-hours per day (between 12:00 and 16:00) to replicate the timescale utilised within the *in vivo* experiments. These conditions were applied to MG63s for 7 and 14 days, and for MSCs for 28 days.

### Quantitative real-time PCR

Subsequently, cells were lysed, RNA extracted and reverse transcription performed using TaqMan Fast Advanced Cells-to-CT Kit (ThermoFisher) according to the manufacturer’s instructions. Forward and reverse primers for qRT-PCR are shown in Table 1. The housekeeping gene used was GAPDH. qRT-PCR was then performed using the QuantStudio 5 Real-Time PCR system (Thermo Fisher Scientific). ΔCt values were calculated using GAPDH and compared between treatment groups. Three biological replicates were used per group, and three technical replicates per sample.

**Table 1.**
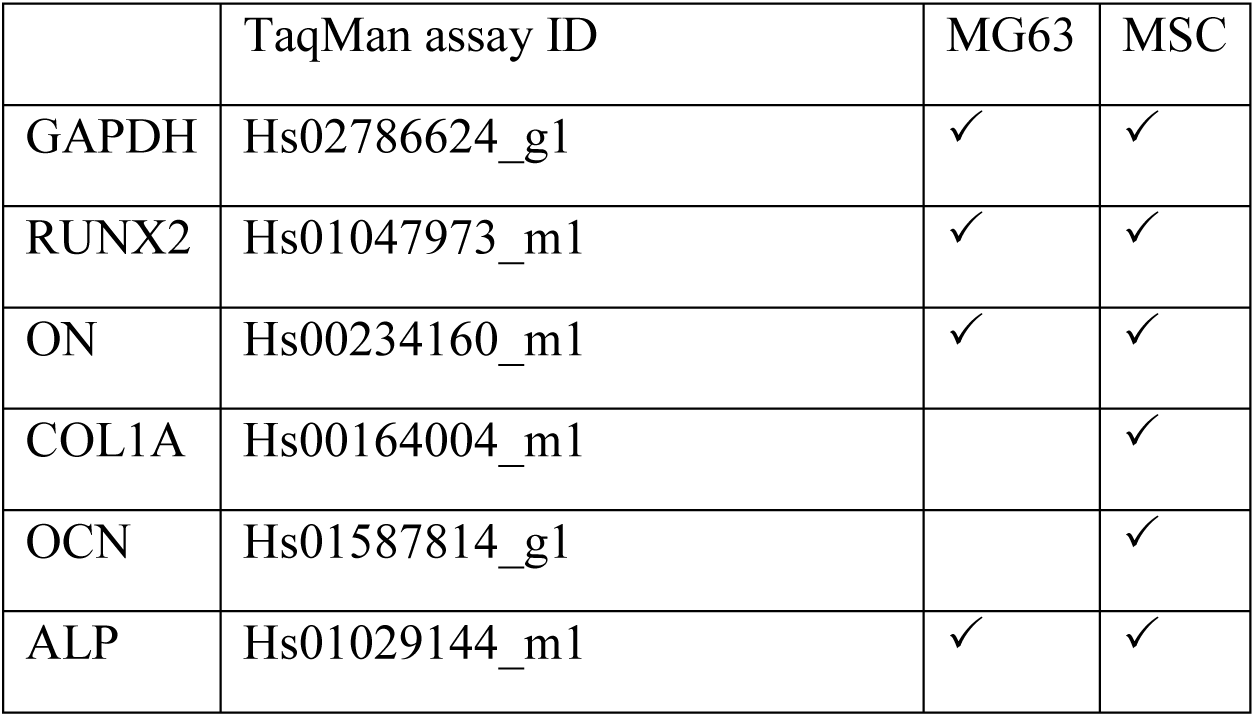
Assay IDs used in qRT-PCR.

### Nanovibration delivery device design

Nanovibration delivery devices and associated electronic systems were designed, manufactured and validated in-house specifically for this study (Figure 1C - F). The device consisted of a bone conduction transducer (Adafruit Industries, New York) and accelerometer (ACH-01, TE Connectivity, Schaffhausen, Switzerland) housed within a custom-made, 3D-printed plastic harness (PLA, 70% infill, resolution 300 µm) (Figure 1C, D). The design featured two holders one for the transducer and accelerometer, respectively. To prevent unwanted vibrations travelling through the device, a strip of foam material (PORON Vive, Algeos, Liverpool) was glued between the two holders. Each holder contained a slot that allowed the passage of an elasticated strap. The strap had hook & loop fastener at its ends, allowing the transducer and accelerometer to remain in firm contact with the lateral and medial sides of the rat hindlimb just below the knee (Figure 1D, E), respectively.

A wave generator circuit was designed, constructed and tested to drive the transducer top plate at 1 kHz, the amplitude of vibration was controlled by the operator with a rotatory potentiometer (See Supplemental Information 1 for further details). Accelerometer circuitry was also designed, constructed and tested to amplify and record the nanoscale vibration detected by the accelerometer. Furthermore, this signal was sent to a Cambridge Electronic Design (CED) Micro 1401 data acquisition unit (CED Limited, Cambridge, UK), connected to a PC, where all the raw data of the measurement session, as well as the average peak value of the signal over each one-minute time scale, were recorded by Spike2 software (associated with CED Limited hardware) on the PC. The Spike2 script also indicated to the operator in real-time whether the acceleration (converted to displacement) measured was within acceptable limits by plotting data coloured red if it was not within the limits and green if it was. These predefined limits were 35 – 45nm, and 90 – 100nm for N40 and N100 groups, respectively. Prior to use in the intervention, accelerometers were calibrated against an *in vitro* nano-amplitude vibration plate, which was itself calibrated using laser interferometry^10^ (See Supplemental Information 2 for further details). The wave generator and accelerometer circuitry were housed in sets of three, giving the capability of nanovibrating multiple rats simultaneously (Figure 1F).

### Interferometric measurement

Measurements of the transmitted vibration amplitude were performed on four SCI rats (3-weeks post-surgery), prior to commencing the intervention to confirm that suitable nanovibration parameters were delivered to hindlimb long bones. Under general anaesthesia, rat hindlimbs were shaved and the device attached. The anteromedial surface of the right proximal tibia and distal femur were surgically exposed. Retroreflective tape was then attached directly to these exposed bone surfaces. Single point laser interferometry (Model SP-S SIOS Meßtechnik GmbH, Ilmenau, Germany) was then performed to measure the amplitude of vibration at 1 kHz from the tape whilst the hindlimb was undergoing direct nanovibration from the device. The amplitude of vibration of the transducer top plate was controlled by the operator with the rotatory potentiometer incrementally increased from the lowest to highest setting. Simultaneously, accelerometer-derived vibration amplitudes from the skin surface just below the exposed proximal tibia were simultaneously acquired (See Figure 2). Multiple measurements were made per rat to observe the expected variation. The spread of nanovibration was also measured along the length of the long bones by exposing other bone sites (mid-femur, distal femur, mid-tibia and distal tibia). Propagation of nanovibration to the contralateral hindlimb was also monitored, by measuring the vibration amplitude at the exposed anteromedial surface of the contralateral proximal tibia.

### Rat model of complete spinal cord transection

Twenty-six male Sprague-Dawley rats weighing 201 – 225g were acquired from Charles River Laboratories (Kent, UK). Rats were housed in threes or fours, in a temperature-controlled room under a 12-hour light-dark cycle, with *ad libitum* access to food and water. All experimental procedures were approved by the Ethical Review Panel of the University of Glasgow and carried out in accordance with the Animals (Scientific Procedures) Act 1986.

Following 1-week of acclimatisation, rats were randomly assigned into two groups: a spinal cord transection (SCT) group (*n* = 20) or sham surgery (SHAM) group (*n* = 6). SCT rats underwent transection of the spinal cord at T9, and in the SHAM group the spinal cord was exposed but not transected. This procedure has been described previously.^12^ Briefly, the spinal cord of anesthetised rats was exposed by laminectomy at the T9-T10 level. The transection was produced by making a small hole in the dura and cutting the spinal cord transversely at two locations, approximately 1 mm apart. The spinal cord tissue between the transections was removed by aspiration and the completeness of the transection confirmed visually through an operating microscope. Rats received buprenorphine (0.05 mg/kg s.c.) and carprofen (5 mg/kg s.c.) the morning of and morning after surgery. Saline (3–5 ml s.c.) and enrofloxacin (5 mg/kg s.c.) were given for 7-days post-surgery. The bladders of SCI rats were manually expressed 3-times per day until spontaneous voiding returned.

Starting 3-days post-surgery, once sufficiently recovered, SCT rats underwent thrice weekly pouch-training sessions for the first 6-weeks post-surgery, to acclimatise to the experimental set up. This involved lightly restraining the rat inside a soft towel pouch, which allowed for access to the hindlimbs (Figure 1E). The length of pouch training sessions was increased weekly from 15-, 30-, 60- to 120-minutes. These training sessions allowed identification of the SCT rats most suitable for undergoing the unilateral nanovibration intervention. After 6-weeks of pouch training the rats suitable for receiving nanovibration were selected. Suitable rats were those that tolerated being pouched for two hours per session. In total ten SCT rats received targeted nanovibrational stimulation. These rats were further subdivided into two nanovibration groups according to vibration amplitude: 40 nm (N40) and 100 nm (N100) groups. SCT rats not selected for vibration were assigned to the spinal cord injury control group (SCI) and rats that received SHAM surgery were assigned to the age-matched control group (AGE-CTR). The four groups that make up this study are; N40 (*n* = 6), N100 (*n* = 4), SCI (*n* = 10) and AGE-CTR (*n* = 6). Three further SCT rats were used to confirm transmission of nanovibration using laser interferometry (as described above).

### Micro-computed tomography

Trabecular and cortical bone morphology and densitometry of the tibia and femur from both hindlimbs were assessed with *ex vivo* micro-Computed Tomography (µCT) using the Bruker SkyScan 1172 scanner (Kontich, Belgium) with Hamamatsu 80 kVp/100 μA X-ray tube at 10 µm isotropic voxel size, as previously described.^40^ All long bones were scanned with the following settings. 70 kVp X-ray tube voltage, 100 μA X-ray tube current, 470 ms exposure time, 2000 × 1332 pixels per image, with a frame averaging of 2, and a 0.4° rotation step for a total of 180° with a 0.5 mm thick aluminium filter. At this voxel size, each long bone was fully captured with either 4 or 5 sub-scans which are stitched together with averaging during reconstruction in NRecon software (Version 1.6.9.18, Kontich, Belgium).

Three volumes of interest (VOIs) were selected for each tibia and femur in CT-Analyser software (Version 1.18.8.0+). For the tibiae, these were the proximal epiphyseal trabecular bone, proximal metaphyseal trabecular bone and mid-diaphyseal cortical bone. For the femora, these were the distal epiphyseal trabecular bone, distal metaphyseal trabecular bone and mid-diaphyseal cortical bone. For epiphyseal trabecular bone, the entire epiphysis enclosed by the growth plate was selected. A percentage-based selection approach was used for the remaining VOIs. The metaphyseal trabecular VOI began at an offset of 2.5% bone length from the growth plate reference point and extended for 5% bone length. The cortical mid-diaphyseal VOIs extended between 47.5% and 52.5% bone length from the proximal end. Epiphyseal trabecular bone was manually segmented from the encapsulating cortical shell. Metaphyseal trabecular and cortical bone VOIs were automatics segmented using a morphological escalation in CT-Analyser, as previously described.^26^

Morphometric analysis was performed on these VOIs after binarization via a global threshold (90/255) and subsequent despeckling for noise removal in CT-Analyser. Trabecular measures included: bone volume fraction (BV/TV), trabecular thickness (Tb.Th), trabecular number (Tb.N), trabecular separation (Tb.Sp) and connectivity density (Conn.D) as per (Bouxsein *et al.*, 2010).^41^ Cortical measures included: cortical thickness (Ct.Th), cortical bone volume (Ct.V), total volume enclosed by the periosteum (Tt.V), marrow volume (Ma.V), cortical volume fraction (Ct.V/Tt.V), second polar moment of area (J), cortical bone surface area to volume ration (BS/BV) as per (Bouxsein *et al.*, 2010) and eccentricity (Ecc).Trabecular volumetric bone mineral density (vBMD) and cortical bone tissue mineral density (TMD) were determined after calibration using two scanner manufacturer provided 4 mm diameter calibration hydroxyapatite phantoms, with known densities of 0.25 and 0.75 g cm^-3^.

Following morphometric analysis in CT-Analyser, the degree of anisotropy (DA) was calculated in BoneJ2^42^ DA is a measure used to quantify the predominant orientation/directionality of trabecular bone. To obtain meaningful values it must be applied to a sample of a larger whole (sub-VOI). DA was determined for a cubic sub-VOI with side length 1.2 mm taken from the proximal tibial metaphyseal trabecular bone VOI. This size of cube was chosen to ensure that the sub-VOI contained at least 5 intra-trabecular lengths.^41^ Consistent placement of the cubic sub-VOI was crucial for obtaining meaningful results, small variation in location would mean that biomechanically homologous regions were not being compared between bones. Consistent placement of the cubic sub-VOI was ensured by spatially aligning all datasets in a semi-automated fashion using a procedure termed co-registration in DataViewer software (Version 1.7.4.2, Kontich, Belgium), as per published methods.^40^ The cubic VOI started 1 mm distal of the proximal tibial growth plate to ensure that i) it only contained secondary spongiosa, and ii) that its location lay within the region directly stimulated by the nanovibration delivery device. The location for the cubic sub-VOI was within the lateral segment of the proximal tibial metaphysis, it was the only region that satisfied all the above requirements (see Supplemental Information 11). The mean intercept length (MIL) algorithm was used to calculate DA.^43^ Briefly, parallel lines from different directions are drawn through the whole cubic sub-VOI. Each individual line is sampled to find points where there is a phase change in the binarized dataset - changes from background to foreground (bone). After all lines are sampled in a given direction, a MIL vector is obtained for that direction, the length of which is equal to the total length of all the lines in that direction divided by the total number of phase changes detected. This is repeated for all the directions. Each MIL vector is then plotted around the origin. An ellipsoid is then fitted to this MIL vector space. It is the radii of the ellipsoid (*a*, *b*; and *c*) that determined DA to be,

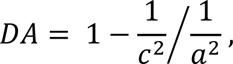

where *a* ≤ *b*; ≤ *c*. The following parameters were used for the analysis; number of directions was set to 2000, lines per direction was set to 10000 and the sampling increment was set to 1.73. The DA algorithm is stochastic because the directions of parallel lines are randomly chosen, this means exactly repeatable results are not guaranteed. The algorithm was run 5 times per sub-VOI to establish the DA. Note that a DA of 0 indicates that the trabecular bone dataset is completely isotropic, while a DA of 1 indicates that it contains a very prominent overall orientation.

Also subsequent to the morphometric analysis, a survey of the 2D trabecular morphometry was conducted along the entire length of each tibia and femur, as per published methods,^40^ to compliment the site-specific 3D trabecular morphometric analysis, and to quantify the regions not typically quantified by that approach, to determine whether there were structural effects of nanovibration that would otherwise have been missed. Briefly, the first 10% bone length proximal and last 10% distal of the tibia, and first 15% proximal and last 15% distal of the femur were excluded from the analysis so to avoid inclusion of the complex geometry of the epiphyses. The remaining trabecular structures were automatically segmented from the cortical bone in CT-Analyser. 2D slice-by-slice analysis of trabecular bone area fraction (BA/TA), was performed in CT-Analyser. BA/TA is defined as the ratio of the total number of pixels representing trabecular bone to the total number of marrow cavity pixels. BA/TA was determined for every single slice in the binarized, segmented trabecular dataset and plotted as a function of bone length. If interesting effects were noticed in specific regions, then these regions of interest could then be further investigated with the standard analysis described above.

### Finite element modelling

A µCT scan of the distal femoral metaphyseal trabecular bone from a representative SHAM group was used to create a 3D model (STL file) in CT-Analyser. This surface mesh was imported into ANSYS SpaceClaim and was cleaned up using the built-in auto-fix function, reducing the number of facets, and shrink-wrapping the body. Multiple iterations of these procedures were required to produce a model that was processable as a volumetric mesh. Harmonic analysis was performed on this mesh to evaluate its structural response when subjected to nanovibration (sinusoidally varying displacement (30 nm, 1 kHz). The material properties of bone were assigned as homogenous, isotropic and linear elastic materials. Specifically, density was acquired from the trabecular bone TMD of the µCT scan, a Young’s modulus of 19.72 GPa, derived from TMD using an empirically-derived equation,^44^ and Poisson’s ratio of 0.3 were used. A displacement was applied to the model in the transverse direction, with a contact area comparable to that of the surface area of the transducer. Elastic supports were assigned to the top and bottom surfaces to simulate adjacent bone structures and a foundational stiffness of 1 N/mm^3^ was used. The model was then subjected to the harmonic analysis.

### Three-point bend mechanical testing

Following μCT scanning, all femora underwent loading to failure in a three-point bend test. Femora were oriented in the anterior-posterior position (with the anterior surface in tension). The actuator head was lowered at a rate of 1 mm min^-1^ using a servohydraulic testing machine with a 2 kN load cell (Zwick/Roell z2.0, August-Nagel-Strasse 11, Ulm, Germany). Femora were preloaded to 10 N and allowed to adapt for 10 seconds before being tested to failure. Load and actuator displacement were recorded at a sampling rate of 100 Hz, using testXpert II (Version 3.61) software. A 15 mm span length was used. The whole-bone structural properties; maximum load, stiffness and absorbed energy were obtained, and the tissue-level mechanical properties; elastic modulus and ultimate stress were calculated from the equations of beam theory.^45^

### Serum bone formation and resorption markers

Serum markers of bone formation and resorption were measured using Rat/Mouse Procollagen type 1 N-terminal propeptide (P1NP) and RatLaps^TM^ C-terminal telopeptide of type I collagen (CTX-I) enzyme immunoassay kits (Immunodiagnostic Systems, Tyne & Wear, UK), respectively, at the time of euthanasia for all rats within the nanovibration study (*n* = 26). The assays were performed following manufacturer’s instructions.

### Metabolomics

Serum samples (*n* = 4 per group, except N100 where n = 3) were defrosted at room temperature for 1.5 hours. Serum metabolites were extracted using a chloroform:methanol:water (1:3:1 ratio) extraction buffer. Samples were vortexed for 5 minutes then centrifuged for 3 min at 13,000g, both at 4.°C. The samples were subsequently used for hydrophilic interaction liquid chromatography-mass spectrometry analysis (Dionex UltiMate 3,000 RSLC, Thermo Fisher, using a 150 × 4.6 mm, 5 µm ZIC-pHILIC column, Merck Sequant). The column was maintained at 40°C and samples were eluted with a linear gradient (20 mM ammonium carbonate in water and acetonitrile) over 26 min at a flow rate of 0.3 ml/min. The injection volume was 10 μl and samples were maintained at 5°C prior to injection. For mass spectrometry analysis, a Thermo Orbitrap QExactive (Thermo Fisher) was used. A standard pipeline, consisting of XCMS54 (peak picking), MzMatch55 (filtering and grouping) and IDEOM56 (further filtering, post-processing and identification) was used to process the raw mass spectrometry data. Identified core metabolites were validated against a panel of unambiguous standards by mass and retention time. Further putative identifications were allotted by mass and predicted retention time. Means and standard errors of the mean were generated for every group of picked peaks and the resulting metabolomics data were uploaded to MetaboAnalyst (Version 5.0)^46^ where heatmap and principal component analyses were performed. Further, the KEGG database and Ingenuity Pathway Analysis software were used to perform an analysis of metabolomic pathways.

### Statistics

To determine differences in gastrocnemius muscle mass, µCT derived morphometric and densitometric parameters, and three-point bend-derived mechanical properties between left and right tibiae and femora within the same group of rats, firstly normality was assessed using the Shapiro-Wilk test on residues and by visually inspecting the spread of data. If data could be assumed normally distributed, then the parametric paired *t*-test was performed. If data could not be assumed normally distributed, then the non-parametric paired samples Wilcoxon test was performed. Multiple group comparisons were performed on right hindlimb tibiae and femora only. Firstly, normality was assessed using the Shapiro-Wilk test on residues and by visually inspecting the spread of data. Homogeneity of variances was tested using Levene’s Test. Data assumed normally distributed and with homogeneity of variances was tested using one-way analysis of variance (ANOVA) with Tukey’s HSD post-hoc. In the cases of normally distributed data but with non-homogeneous variances, ANOVA was performed with Games Howell post hoc test, while data assumed to not be normally distributed data was tested with independent samples Kruskal-Wallis test for multiple groups with Dunn’s post-hoc test. To determine differences between left and right tibiae (and left and right femora) within the same group of rats, firstly normality was assessed using the Shapiro-Wilk test on residues and by visually inspecting the spread of data. If data could be assumed normally distributed, then the parametric paired *t*-test was performed. If data could not be assumed normally distributed, then the non-parametric paired samples Wilcoxon test was performed. A mixed-model repeated measures ANOVA was used to assess body mass at multiple time points within the same rats. Significance was defined as *p* < 0.05. For qRT-PCR data, Dixon’s Q test for outliers was performed with significance level set to *p* < 0.15, subsequently one-way ANOVA with Tukey’s HSD post-hoc was performed. All results are expressed as mean ± standard deviation. All statistical analyses were performed in R (Version 3.6.1).

## ASSOCIATED CONTENT

### Supporting Information Available

Description of the nanovibration delivery device electronic systems, calibration of accelerometers, an overview of a preliminary study investigating nanovibration as an intervention for the prevention/attenuation of SCI-induced osteoporosis. Figures of initial and end rat body mass, rat body mass throughout intervention, gastrocnemius muscle mass at end of intervention. Figures of µCT analysis of proximal tibial metaphyseal and epiphyseal trabecular bone, distal femoral metaphyseal and epiphyseal trabecular bone and three-point bend-determined whole-bone and material-level mechanical properties of tibial mid-diaphyseal cortical bone. Schematic of location of trabecular bone volume of interest for degree of anisotropy analysis (DOC). Data collected in this study is accessible at “”. µCT datasets can be made available on reasonable request.

### Corresponding Authors

Jonathan A. Williams - Department of Biomedical Engineering, Wolfson Building, University of Strathclyde, Glasgow, G4 0NW, United Kingdom &. Institute of Neuroscience and Psychology, College of Medical, Veterinary and Life Sciences, University of Glasgow, G12 8QQ, United Kingdom & Scottish Centre for Innovation in Spinal Cord Injury, Queen Elizabeth National Spinal Injuries Unit, Glasgow, United Kingdom;

Stuart Reid - Department of Biomedical Engineering, Wolfson Building, University of Strathclyde, Glasgow, G4 0NW, United Kingdom;

### Author Contributions

J.A.W, S.C., M.J.D, J.S.R and S.R. conceived the experiments. J.A.W., P.C., R.G., O.J-L, A.W., M.S., R.M., C.H., and J.S.R. performed the experiments. J.A.W., P.C., P.G.C, C.H., J.F.C.W., M.P., M.J.D. and P.C. provided materials and expertise. M.P., S.C., J.S.R. and S.R. acquired funding. J.A.W, O.J-L, A.W., and R.M. analysed the data. J.A.W. wrote the manuscript and prepared the figures. All authors have given approval to the final version of the manuscript.

## ACKNOWLEDGEMENT

This study was supported by funding provided by the Science and Technology Facilities Council (STFC) Challenge Led Applied Systems Programme (ST/S000968/1) and (ST/S000852/1), and support from the Universities of Strathclyde and Glasgow. R.G. was supported by the Engineering and Physical Sciences Research Council (EPSRC) Centre for Doctoral Training in Medical Devices & Health Technologies (EP/L015595/1). S.R. is supported by a Royal Society Industry Fellowship (INF\R1\201072). Data collected in this study is accessible at https://doi.org/10.15129/0b2d2d47-f4c9-40a6-a2ad-72565a740e37

