## Supplemental Information for "Developing and Investigating a Nanovibration Intervention for the Prevention/Reversal of Bone Loss Following Spinal Cord Injury"

| Table of content | | |
|---|---|---|
| **Item** | **Title** | **Page** |


| Item | Title | Page |
|---|---|---|
| **SI1** | Description of nanovibration delivery device electronic systems | **4-7** |
| **SI2** | Calibration of accelerometers | **8-10** |
| **SI3** | Preliminary study investigating nanovibration as an intervention for the prevention/attenuation of SCI-induced osteoporosis | **11-14** |
| **SI4** | Initial and end rat body mass | **15** |
| **SI5** | Rat body mass throughout intervention | **16** |
| **SI6** | Gastrocnemius muscle mass at end of intervention | **17** |
| **SI7** | µCT analysis of proximal tibial metaphyseal trabecular bone – contralateral comparisons | **18-19** |
| **SI8** | µCT analyses of proximal tibial epiphyseal and distal femoral metaphyseal and epiphyseal trabecular bone | **20-28** |
| **SI9** | µCT analysis of tibial mid-diaphyseal cortical bone morphometry | **29-31** |
| **SI10** | Three-point bend-determined whole-bone and material-level mechanical properties of tibial mid-diaphyseal cortical bone | **32** |
| **SI11** | Location of trabecular bone volume of interest for degree of anisotropy analysis | **33** |

**Supplemental Information 1 - Description of Nanovibration Delivery Device Electronic Systems**

The electronics for the animal device consist of two main elements; the wave generator circuit and the accelerometer amplification circuit. The wave generator circuit is based on that used to generate a sine wave voltage signal in the Nanokick bioreactor (Campsie et al., 2019). An AD9833 low power, programmable waveform generator (Analog Devices, Massachusetts, USA) produces the required sine wave with the output frequency and phase programmed using an ATMega328 microcontroller (Atmel, California, USA). Filtering is required at the output of the AD9833 to significantly reduce higher frequency components, created during the digital synthesis of the primary signal, using a seventh order LC elliptical reconstruction filter. A non-inverting amplifier circuit, using an OPA37 ultra-low noise OP-AMP (Texas Instruments, Texas, USA), is utilised to boost the amplitude of the filtered sine wave before it reaches the final amplification stage. A 10 kΩ rotary potentiometer was used for the N40 rats in the non-inverting amplifier circuit allows the gain, and therefore the amplitude of the sine wave to be adjusted by the user. This was switched to a 20 kΩ potentiometer for the N100 rats to allow higher gain, and therefore higher displacement amplitudes from the transducer.

$$GAIN = 1 + \frac{R_{POT}}{R_2} \qquad \text{Equation 1}$$

Equation 1 shows the gain for a non-inverting op-amp. Increasing the maximum value of the potentiometer ($R_{POT}$), which is acting as the feedback resistor in our circuit, increases the gain of the amplifier.

Finally, the sine wave signal is amplified with a MAX98306 3.7W power amplifier (Maxim Integrated, California, USA) to provide the bone conduction transducer with the voltage and current needed to function. The power amplification stage was purchased as a standalone printed circuit board (PCB) (Adafruit Industries, New York, USA). A block diagram of circuitry described above can be seen in Figure 1 and the wave generator and power amplifier PCBs can be seen in Figure 2.

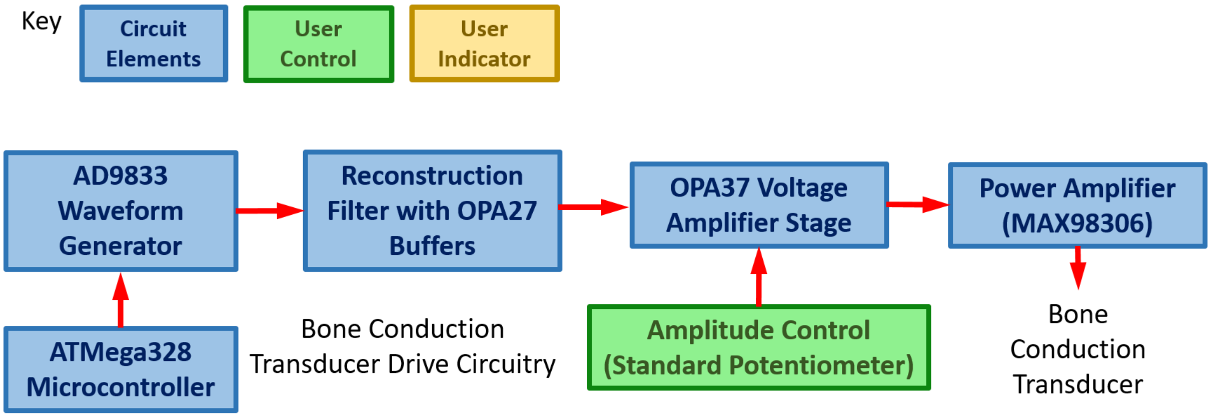


**Figure 1:** Block diagram of main components of the wave generator PCB used in the rat study.

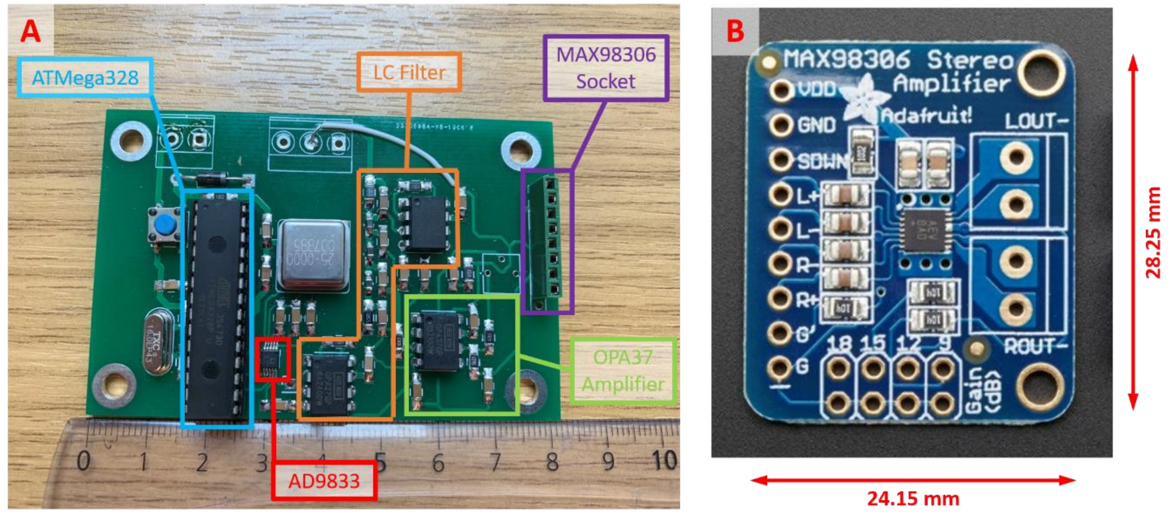


**Figure 2:** (A) Photograph of the wave generator PCB with main components highlighted (B) Image of power amplifier (MAX98306) PCB purchased from Adafruit Industries.

The vibration from the bone conduction transducer is detected using an ACH-01 accelerometer (TE Connectivity, Schaffhausen, Switzerland). This particular accelerometer utilises an inertial mass on a piezoelectric polymer film to generate a voltage when an acceleration is detected. Since the accelerations being measured in this instance are at the nanoscale, a low noise, multi-stage amplifier is needed to significantly boost the voltage generated from the ACH-01 device. The first stage of the multi-stage amplification process is a non-inverting amplifier, using an OPA37 OP-AMP, the design of which is given in the ACH-01 data sheet. The second stage is a standard inverting amplifier, also consisting of an OPA37 OP-AMP. The amplified signal is sent to a Cambridge Electronic Design (CED) Micro 1401 data acquisition unit (CED Limited, Cambridge, UK), connected to a PC, where all the raw data of the measurement session, as well as the average peak value of the signal over a one-minute time scale, are recorded by Spike2 software (associated with CED Limited hardware) on the PC. The Spike2 script also indicates to the operator in real time whether the acceleration measured is within acceptable limits for the animal experiments by plotting data coloured red if it is not within the limits and green if it is. A block diagram of this setup is shown in Figure 3.

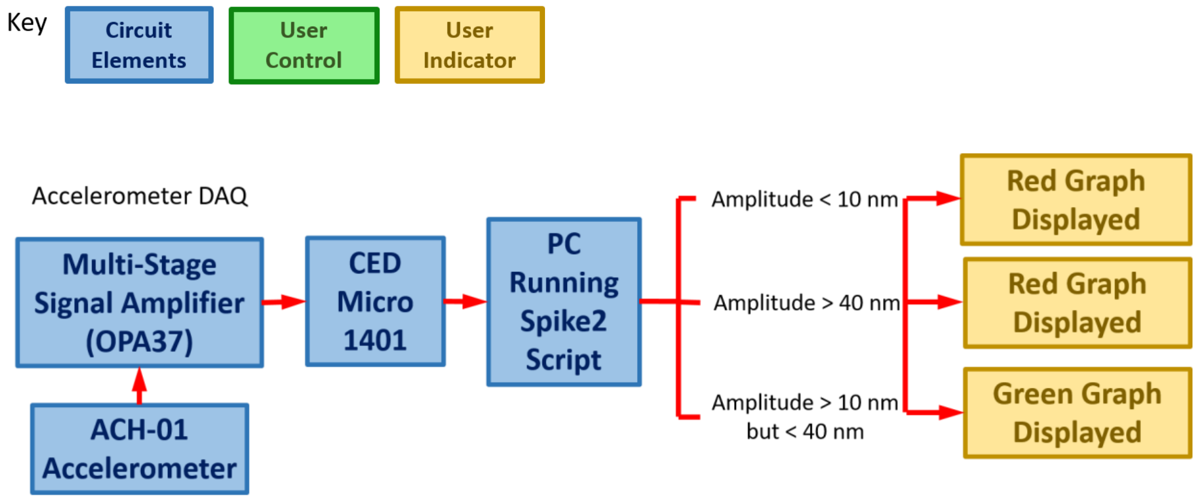

**Figure 3:** Block diagram of main components of the accelerometer circuitry and data acquisition (DAQ) process used in the animal study.

### Supplemental Information 2 – Calibration of Accelerometers

Each of the ACH-01 accelerometers gives a different voltage output for a given acceleration due to differences in the manufacturing process of each individual device. Therefore, all accelerometers used in the studies described here are individually calibrated and have their own unique calibration curve. It is also important to distinguish that in previous nanokicking research the displacement, not acceleration, is the factor that is quantified, therefore, the calibration process allows the device to be calibrated in terms of displacement so that is comparable to previous literature. The calibration process is carried out by magnetically fixing the accelerometer to a Nanokick bioreactor, which creates a precise nanoscale oscillation at 1 kHz and different set amplitudes and measuring the displacement of the accelerometer with a laser interferometer (Model SP-S SIOS Meßtechnik GmbH, Ilmenau, Germany). The Nanokick bioreactor outputs a very precise and stable nanoscale vibration, using an array of piezoelectric ceramics, for cell culture experiments and is itself calibrated by laser interferometry, an instrument that can measure displacements with sub-nanoscale resolution. Before calibration, each accelerometer is individually numbered using a scribe on the plastic casing so it is easily identified, a small section of self-adhesive rubber magnet is fixed to the underside of the accelerometer to attach it to the bioreactor top plate, and retroreflective tape is attached to the top surface of the accelerometer to reflect as much of the interferometer's laser light back to its' sensor as possible for an accurate measurement. The bioreactor is powered with a sine wave signal generated from an AFG-21005 arbitrary function generator (GW Instek, New Taipei City, Taiwan) and amplified by a Behringer KM750 amplifier (Behringer, Willich, Germany). The frequency is fixed at 1kHz and the displacement amplitude is adjusted by changing the output voltage amplitude on the function generator. Two sets of calibration measurements were taken because it was found that over a measurement range of 1 – 200 nm the gradient of the calibration curve can be weighted by data points at the higher end of the scale, giving conversions at the lower end of the scale a larger error. For wide range measurements, the amplitude of the sine wave is incrementally increased from 2.5 nm to 200 nm and the ACH-01 voltage measured with an oscilloscope ($V_{rms}$ and $V_{peak\text{-}to\text{-}peak}$), see Figure 4B, and for shorter range measurements data is taken from 1 – 10 nm in incremental steps of 1 nm, see Figure 4A. For the displacements detected during animal experiments it was determined that the calibration curve for the lower range measurements would give the most accurate results.

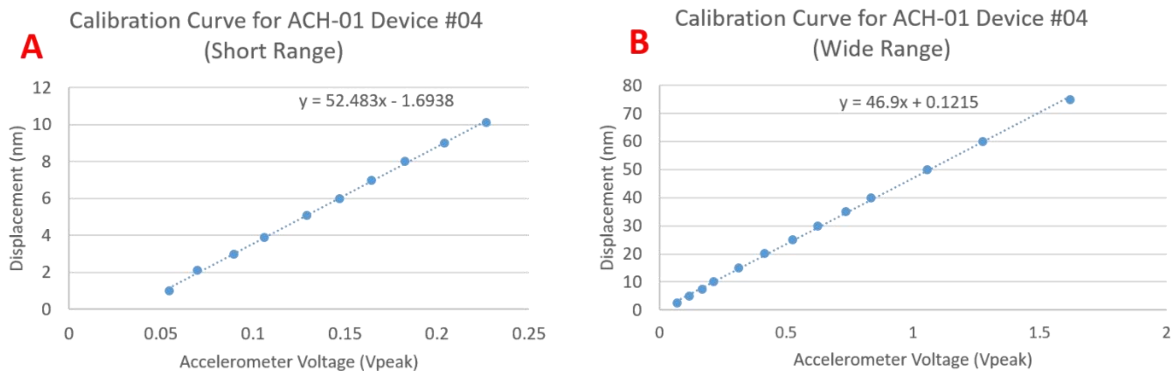


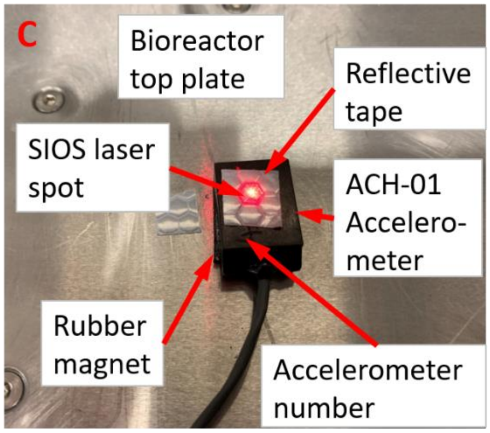


D

| Displacement measured with interferometer (nm) | Displacement measured with calibrated ACH-01 Device #04 (nm) |
|---|---|
| 1.1 | 0.4 |
| 2.5 | 1.8 |
| 5 | 4.4 |
| 7.5 | 7.8 |
| 10 | 10.6 |
| 14.9 | 15.3 |
| 20 | 21.3 |
| 30 | 32.6 |

**Figure 4**: (A) Plot of the calibration data for ACH-01 device 4 over a short range of displacement values (B) Plot of the calibration data for ACH-01 device 4 at the higher range of displacement values (C) Picture of ACH-01 device 4 on bioreactor top plate being measured with the laser

interferometer (D) Comparison of displacement measured with interferometer with data recorded by Spike2 software from calibrated ACH-01 device 4.

During the rat experiments two instrument boxes containing three wave generator PCBs each and two instrument boxes containing three accelerometer amplifier PCBs each were used so that a maximum of six animals per session can receive the nanovibrational intervention.

**Supplementary Information 3: Preliminary study investigating nanovibration as an intervention for the prevention/attenuation of SCI-induced osteoporosis**

This preliminary study was initial setup to investigate the suitability of the designed device. After the devices were assessed and shown to be operating as planned, it was continued for a full 6 weeks to investigate whether nanovibration can slow or prevent the osteoporotic changes that occur after spinal cord transection. This was first time that the proposed nanovibration intervention was been explored *in vivo*.

Briefly, four male Sprague-Dawley rats were acquired from Charles Rivers Laboratories, Kent, UK. Rats were given T9-transections as described in the main manuscript. The nanovibration intervention was started as soon as practicably possible after surgery, which was 3-days. Three of these rats underwent the nanovibration protocol for 6-weeks as described in the main manuscript.

Throughout the study body mass was tracked, end point bone and gastrocnemius muscle masses were measured and μCT scanning and morphometric analysis was performed, where nanovibrated (right) and non-vibrated contralateral control (left) hindlimb were compared.

*Body Mass Results*

Body mass at time of surgery was similar between rats (233 – 245 grams). Due to growth in the rat model between time of surgery and the end of intervention (44-days later) there was a 37% to 63% increase in body mass ($p < 0.001$) (Figure A1). It took between 4-and 12-days for all rats to regain their pre-surgery body mass. The four day turn around point between losing and gaining body mass should not be associated with the nanovibration, this is typical for the rat model.

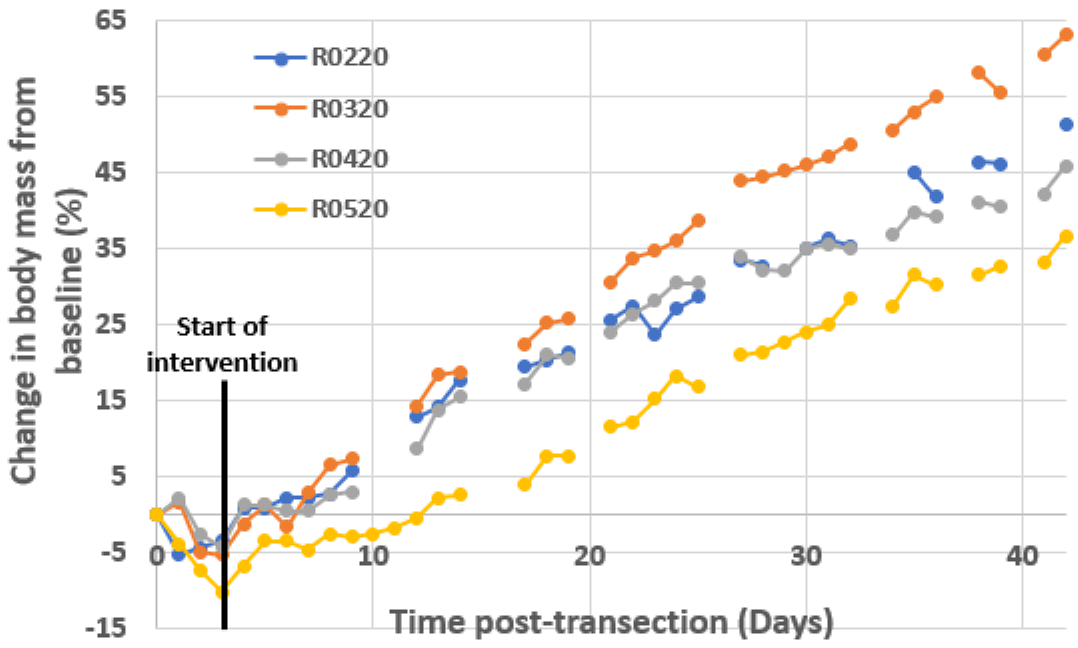


**Figure A1.** Body mass with time post-T9 transection of the spinal cord surgery for rats that underwent the preliminary nanovibration study, starting 3 days post-surgery.

*Macroscopic properties of hindlimb muscle and bone*

No significant difference was observed between right (nanovibrated) and left (control) hindlimb gastrocnemius muscle wet mass (Table A1). Although for two rats the control gastrocnemius mass was higher in the control hindlimb. Also, there was no significant difference in tibial or femoral wet mass, despite in all instance in individual rat left-right comparisons, the left was always heavier.

**Table A1.** Vibrated and control hindlimb muscle and bone wet mass after spinal transection and 6-weeks of nanovibration.

| | Gastrocnemius Mass (g) | | | Tibia mass (g) | | | Femur mass (g) | | |
|---|---|---|---|---|---|---|---|---|---|
| **Rat ID** | **Left** | **Right** | **Diff (%)** | **Left** | **Right** | **Diff (%)** | **Left** | **Right** | **Diff (%)** |
| **R0320** | 2.32 | 1.98 | -14.6 | 0.87 | 0.84 | -2.9 | 0.98 | 0.93 | -5.1 |
| **R0420** | 1.70 | 1.78 | 4.8 | 0.78 | 0.76 | -1.9 | 0.87 | 0.87 | -0.5 |
| **R0520** | 1.96 | 1.56 | -20.1 | 0.74 | 0.66 | -10.5 | 0.86 | 0.79 | -7.9 |
| **Average** | **1.99** | **1.77** | **-10.0** | **0.80** | **0.76** | **-5.1** | **0.90** | **0.86** | **-4.5** |
| **SD** | **0.32** | **0.21** | | **0.06** | **0.09** | | **0.07** | **0.07** | |
| **p-value** | **0.3774** | | | **0.5626** | | | **0.4981** | | |

*Trabecular bone 3D morphometric results*

Representative proximal tibia metaphyseal secondary spongiosa VOIs and morphometric measures are shown in Figure A3 and Table A2. Paired T-tests indicated that there was no significant difference in the volume fraction of trabecular bone (BV/TV) between vibrated and control proximal tibial metaphyseal secondary spongiosa VOIs, or that there was any significant difference between their underlying trabecular microarchitectures. Despite this, BV/TV, trabecular thickness (Tb.Th) and trabecular number (Tb.N) were lower in the vibrated compared to control for all three rats. The same is true for proximal tibia epiphyseal trabecular VOI (Figure A4 & Table A3). A similar trend was observed for the distal femoral metaphyseal and epiphyseal trabecular bone (data not shown)

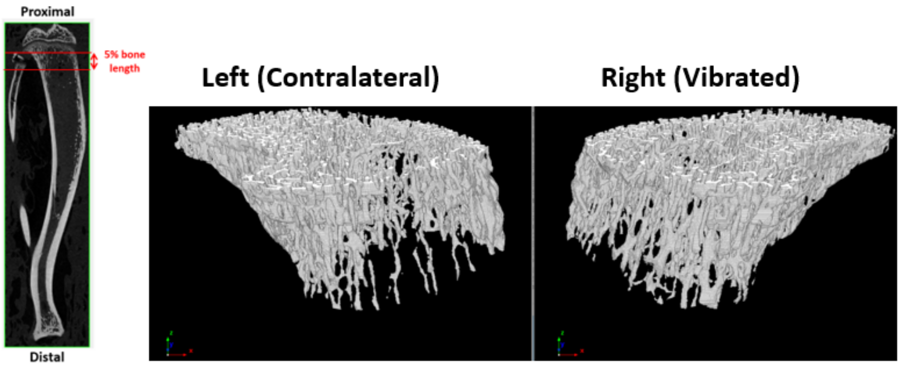


**Figure A3.** Representative μCT-based images of 5% bone length metaphyseal trabecular VOI for vibrated and control hindlimbs.

**Table A2.** Proximal tibial metaphyseal trabecular bone morphometric parameters. Data shown as average ± SD.

| | BV/TV (%) | | | Tb.Th (mm) | | | Tb.Sp (mm) | | | Tb.N (mm$^{-1}$) | | | Conn.D (mm$^{-3}$) | | |
|---|---|---|---|---|---|---|---|---|---|---|---|---|---|---|---|
| **Rat ID** | **Left** | **Right** | **Diff (%)** | **Left** | **Right** | **Diff (%)** | **Left** | **Right** | **Diff (%)** | **Left** | **Right** | **Diff (%)** | **Left** | **Right** | **Diff (%)** |
| **R0320** | 11.92 | 10.34 | -13.3 | 0.071 | 0.069 | -4.1 | 0.49 | 0.54 | -10.4 | 1.67 | 1.51 | -9.5 | 56.3 | 54.1 | -3.9 |
| **R0420** | 8.11 | 7.76 | -4.3 | 0.072 | 0.070 | -3.4 | 0.66 | 0.59 | 10.5 | 1.13 | 1.11 | -1.0 | 30.7 | 33.8 | 10.0 |
| **R0520** | 12.96 | 7.87 | -39.3 | 0.072 | 0.066 | -8.8 | 0.41 | 0.72 | -74.4 | 1.79 | 1.19 | -33.4 | 64.3 | 45.7 | -29.0 |
| **Average** | **11.00** | **8.65** | **-19.0** | **0.07** | **0.07** | **-5.5** | **0.52** | **0.62** | **-24.7** | **1.53** | **1.27** | **-14.6** | **50.4** | **44.5** | **-7.6** |
| **SD** | **2.55** | **1.46** | | **0.00** | **0.00** | | **0.13** | **0.09** | | **0.35** | **0.21** | | **17.6** | **10.2** | |
| **p-value** | **0.2405** | | | **0.0869** | | | **0.4781** | | | **0.2831** | | | **0.4604** | | |

BV/TV: trabecular bone volume fraction, Tb.Th: trabecular thickness, Tb.Sp: trabecular separation, Tb.N: trabecular number, Conn.D: Connectivity Density.

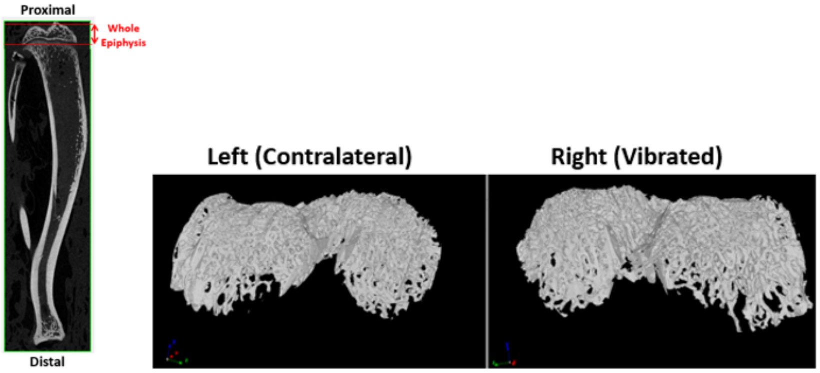


**Figure A4.** Representative μCT-based images of epiphyseal trabecular VOI for vibrated and control hindlimbs.

**Table A3.** Proximal tibial epiphyseal trabecular bone morphometric parameters. Data shown as average ± SD.

| | BV/TV (%) | | | Tb.Th (mm) | | | Tb.Sp (mm) | | | Tb.N (mm$^{-1}$) | | | Conn.D (mm$^{-3}$) | | |
|---|---|---|---|---|---|---|---|---|---|---|---|---|---|---|---|
| **Rat ID** | **Left** | **Right** | **Diff (%)** | **Left** | **Right** | **Diff (%)** | **Left** | **Right** | **Diff (%)** | **Left** | **Right** | **Diff (%)** | **Left** | **Right** | **Diff (%)** |
| **R0320** | 30.34 | 27.15 | -10.5 | 0.102 | 0.096 | -6.3 | 0.24 | 0.25 | 5.8 | 2.97 | 2.84 | -4.5 | 128.7 | 128.5 | -0.2 |
| **R0420** | 26.34 | 25.21 | -4.3 | 0.109 | 0.108 | -0.6 | 0.31 | 0.30 | -2.3 | 2.42 | 2.33 | -3.7 | 66.4 | 71.2 | 7.3 |
| **R0520** | 30.49 | 22.46 | -26.3 | 0.110 | 0.099 | -10.1 | 0.28 | 0.33 | 19.1 | 2.78 | 2.28 | -18.0 | 104.8 | 83.9 | -20.0 |
| **Average** | **29.05** | **24.94** | **-13.7** | **0.11** | **0.10** | **-5.7** | **0.27** | **0.29** | **7.5** | **2.72** | **2.48** | **-8.8** | **100.0** | **94.5** | **-4.3** |
| **SD** | **2.35** | **2.35** | | **0.01** | **0.00** | | **0.03** | **0.04** | | **0.28** | **0.31** | | **31.5** | **30.1** | |
| **p-value** | **0.1816** | | | **0.1830** | | | **0.3746** | | | **0.2054** | | | **0.5601** | | |

BV/TV: trabecular bone volume fraction, Tb.Th: trabecular thickness, Tb.Sp: trabecular separation, Tb.N: trabecular number, Conn.D: Connectivity Density

**Supplementary Information 4: Initial and end rat body mass**

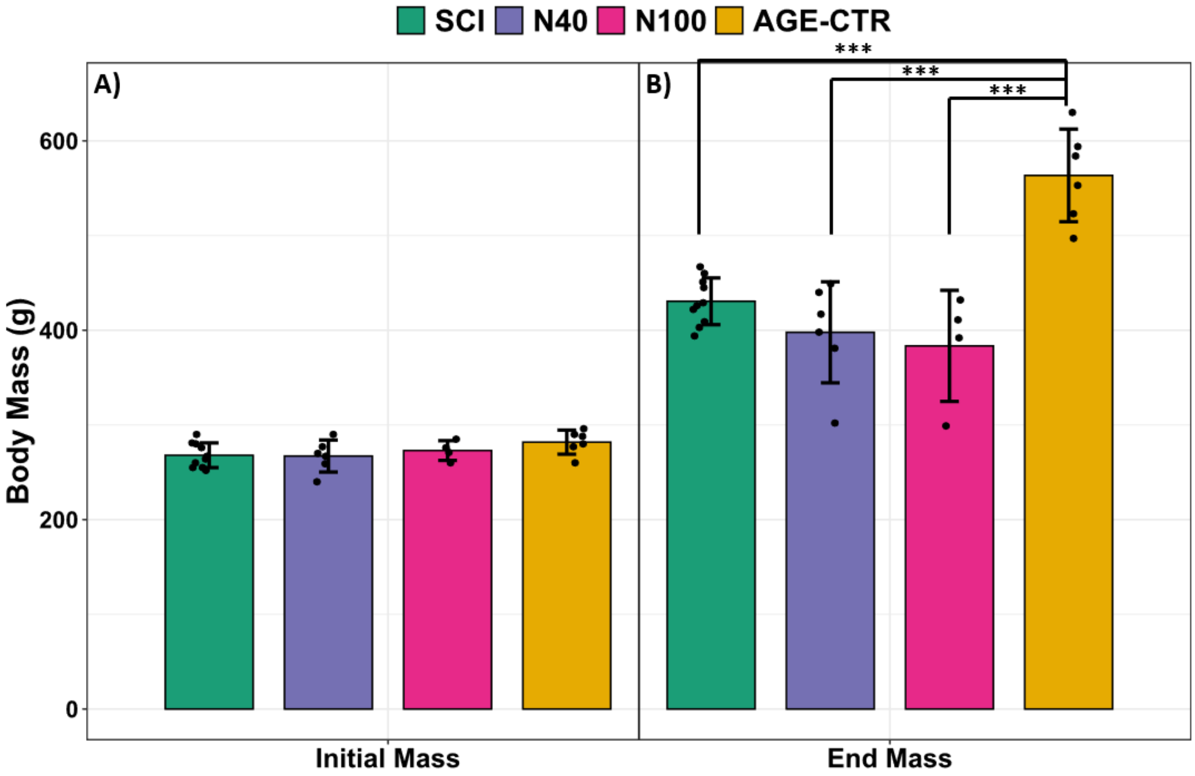


**Figure S4:** Plots of A) initial and B) end body mass of rats for all groups. Data shown as mean ± SD. *** indicates $p < 0.001$.

**Supplementary Information 5: Rat body mass throughout intervention**

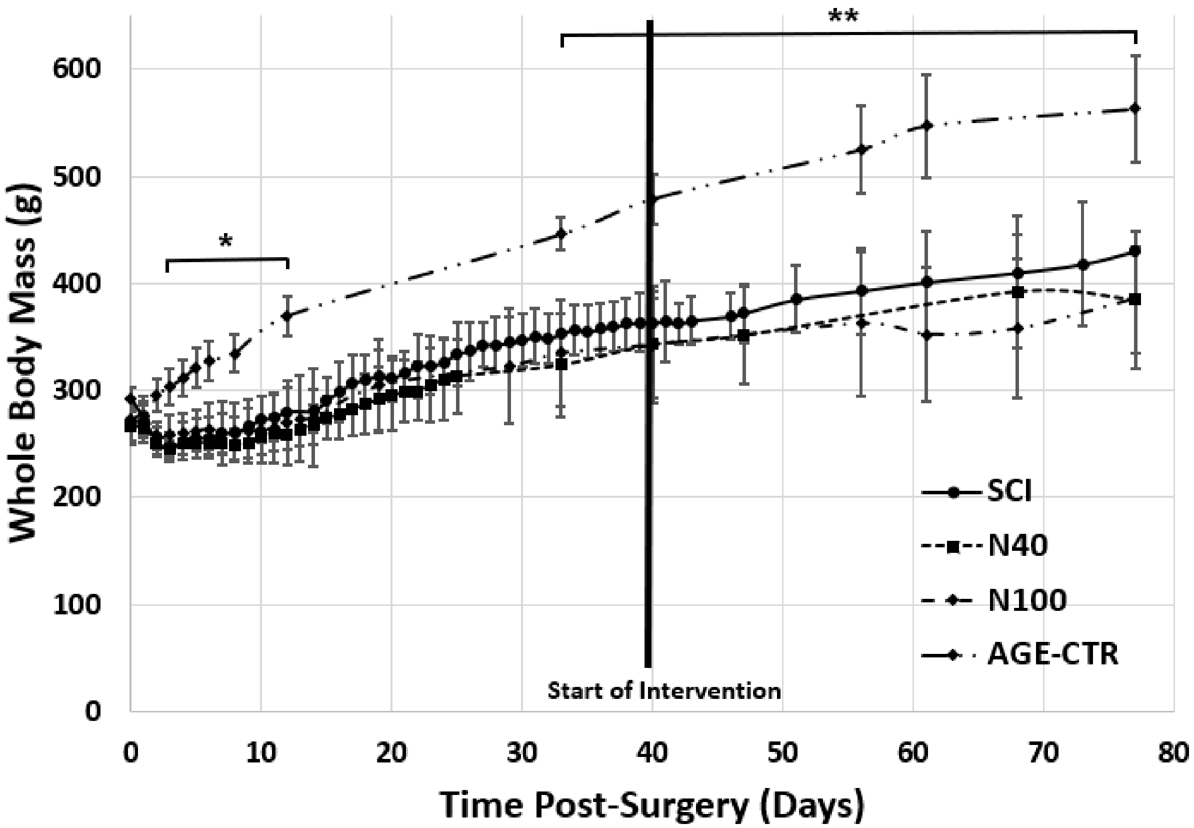


**Figure S5:** Body mass with time post-surgery for all groups. Data shown as mean ± SD. * and ** indicate $p < 0.05$ and $p < 0.01$, respectively, for AGE-CTR versus all other groups at the same post-surgical timepoint. Bold line indicates that 40 to 42 days after surgery N40 and N100 groups start intervention.

**Supplementary Information 6: Gastrocnemius muscle mass**

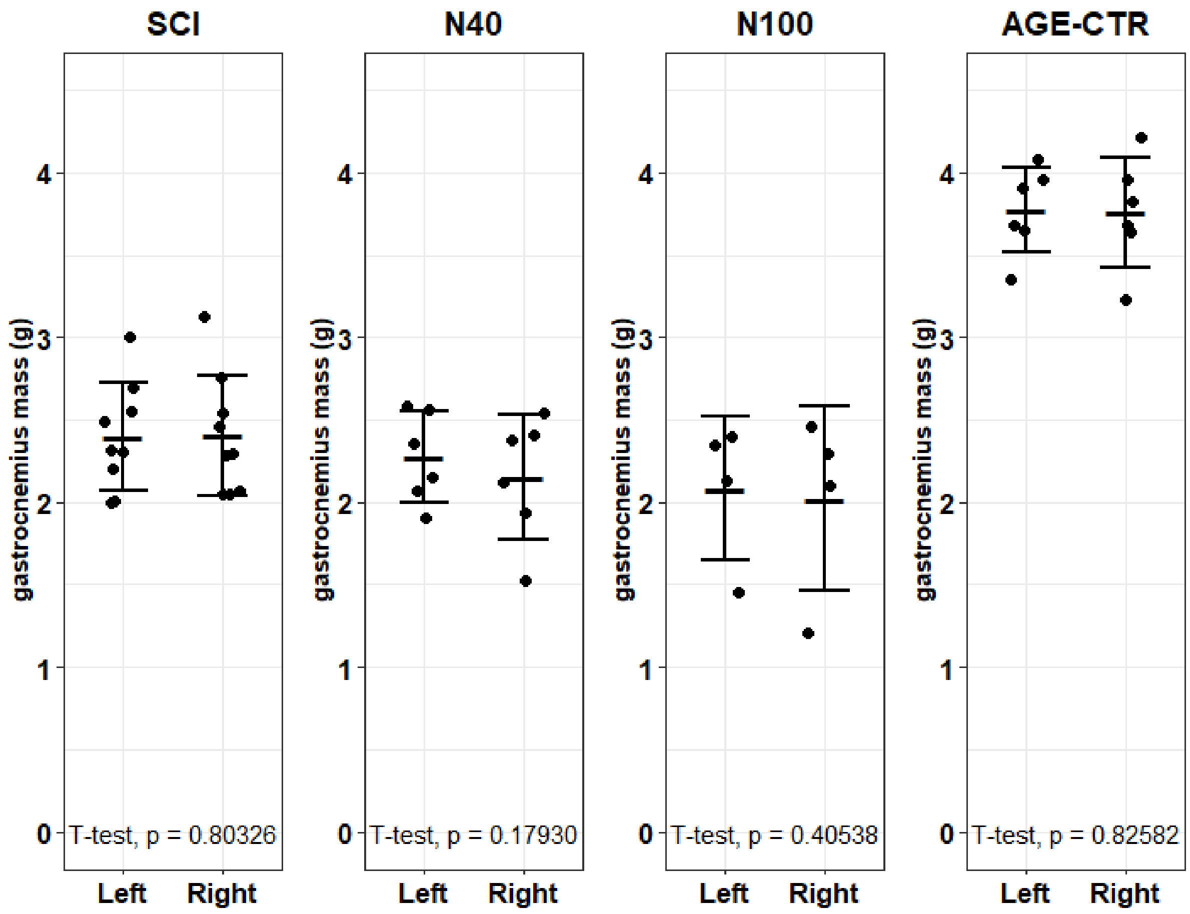


**Figure S6:** Plots of left (contralateral) and right (nanovibrated) hindlimb gastrocnemius muscle mass for all 4 experimental groups. Data shown as mean ± SD.

### Supplementary Information 7: µCT Analysis of proximal tibial metaphyseal trabecular bone – contralateral comparisons

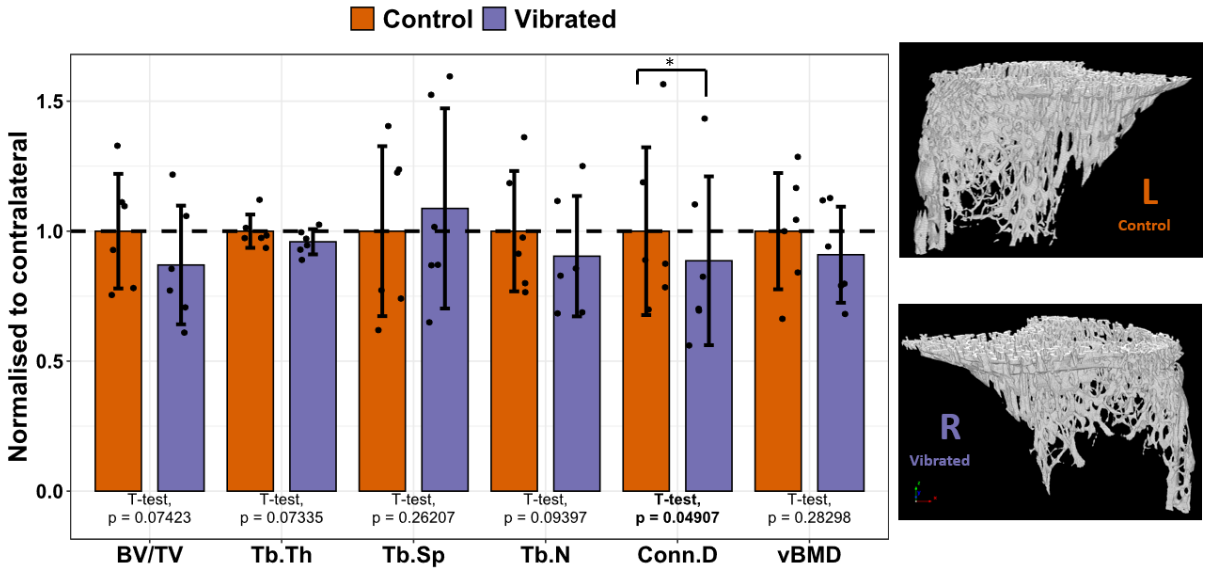


**Figure 7A.** Representative µCT-based images of the nanovibrated (right) and contralateral control (left) proximal tibia metaphyseal secondary spongiosa VOI with mean morphometric outcome measures for 40 nm amplitude vibrated (N40) rats. Data shown as mean ± SD with each parameter normalised to that of contralateral control. * indicates $p < 0.05$.

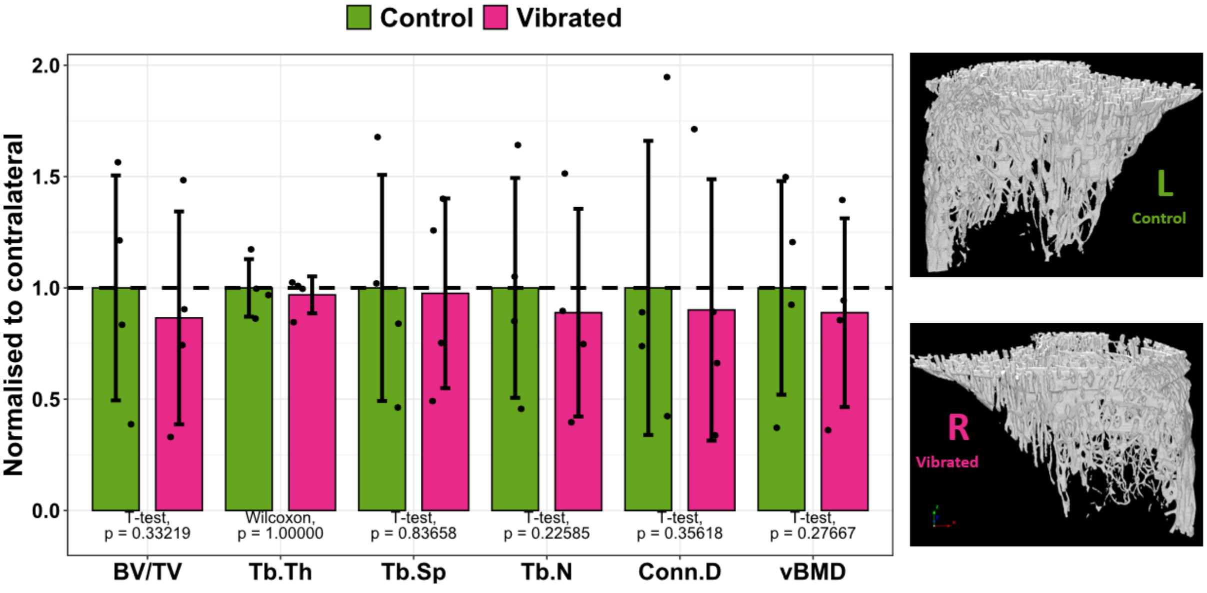

**Figure 7B.** Representative µCT-based images of the nanovibrated (right) and contralateral control (left) proximal tibia metaphyseal secondary spongiosa VOI with mean morphometric outcome measures for 100 nm amplitude vibrated (N100) rats. Data shown as mean ± SD with each parameter normalised to that of contralateral control.

**Supplementary Information 8: μCT Analyses of proximal tibial epiphyseal and distal femoral metaphyseal and epiphyseal trabecular bone**

*i)* *Proximal tibial epiphyseal trabecular bone*

- *Contralateral comparison of N40 group*

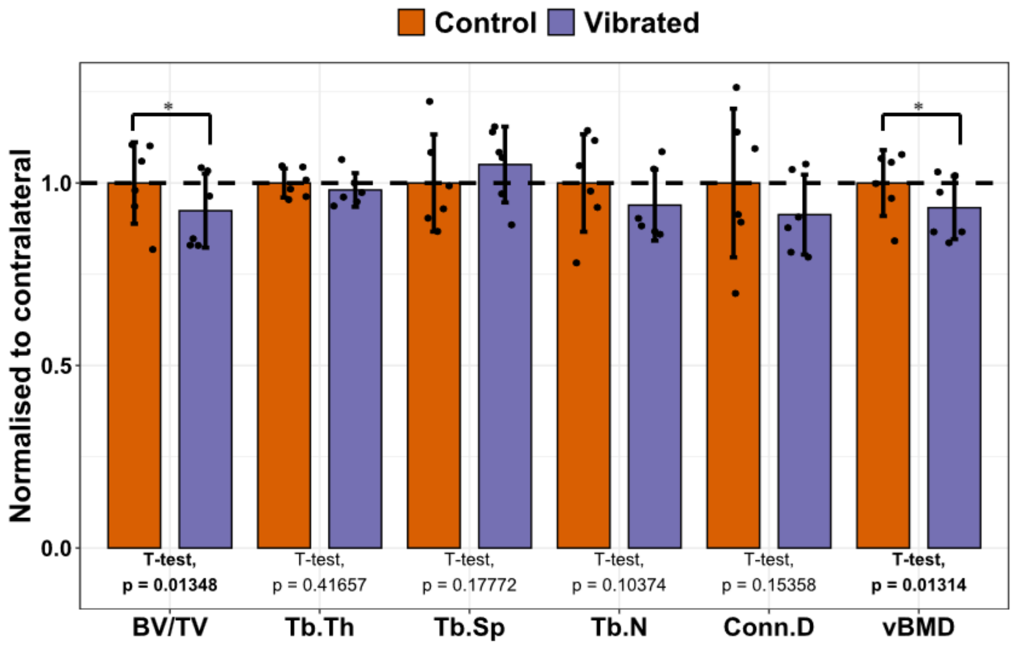


**Figure 8A**. Mean morphometric outcome measures of the nanovibrated (right) and contralateral control (left) proximal tibial epiphyseal secondary spongiosa VOI for 40 nm amplitude vibrated (N40) rats. Data shown as mean ± SD with each parameter normalised to that of contralateral control. * and ** indicate $p < 0.05$ and $p < 0.01$, respectively.

- *Contralateral comparison of N100 group*

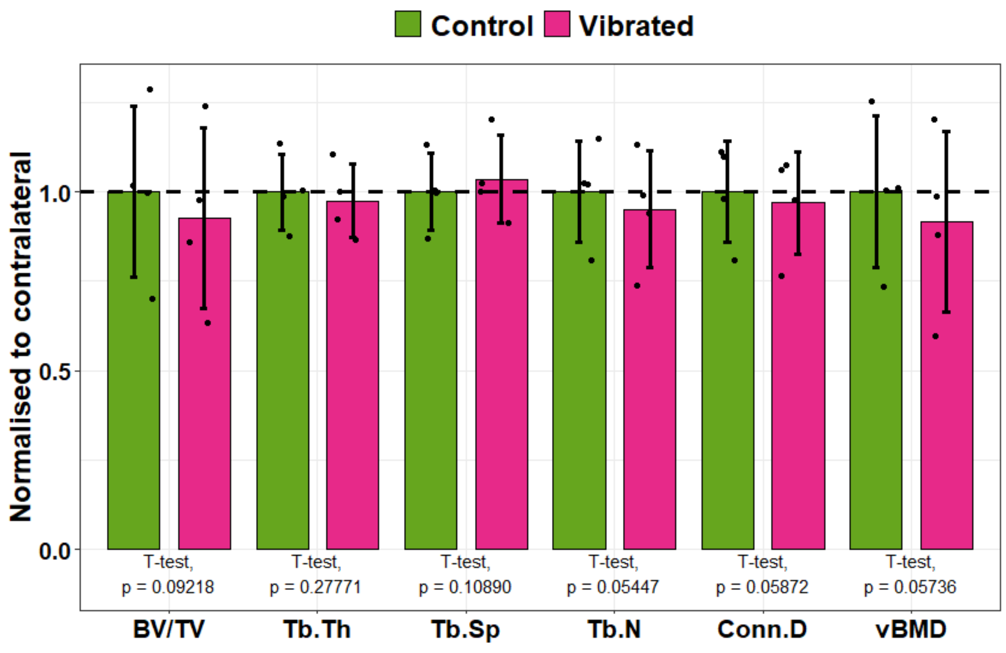


**Figure 8B**. Mean morphometric outcome measures of the nanovibrated (right) and contralateral control (left) proximal tibial epiphyseal secondary spongiosa VOI for 100 nm amplitude vibrated (N10) rats. Data shown as mean ± SD with each parameter normalised to that of contralateral control.

- *Comparison of nanovibrated/right between all groups*

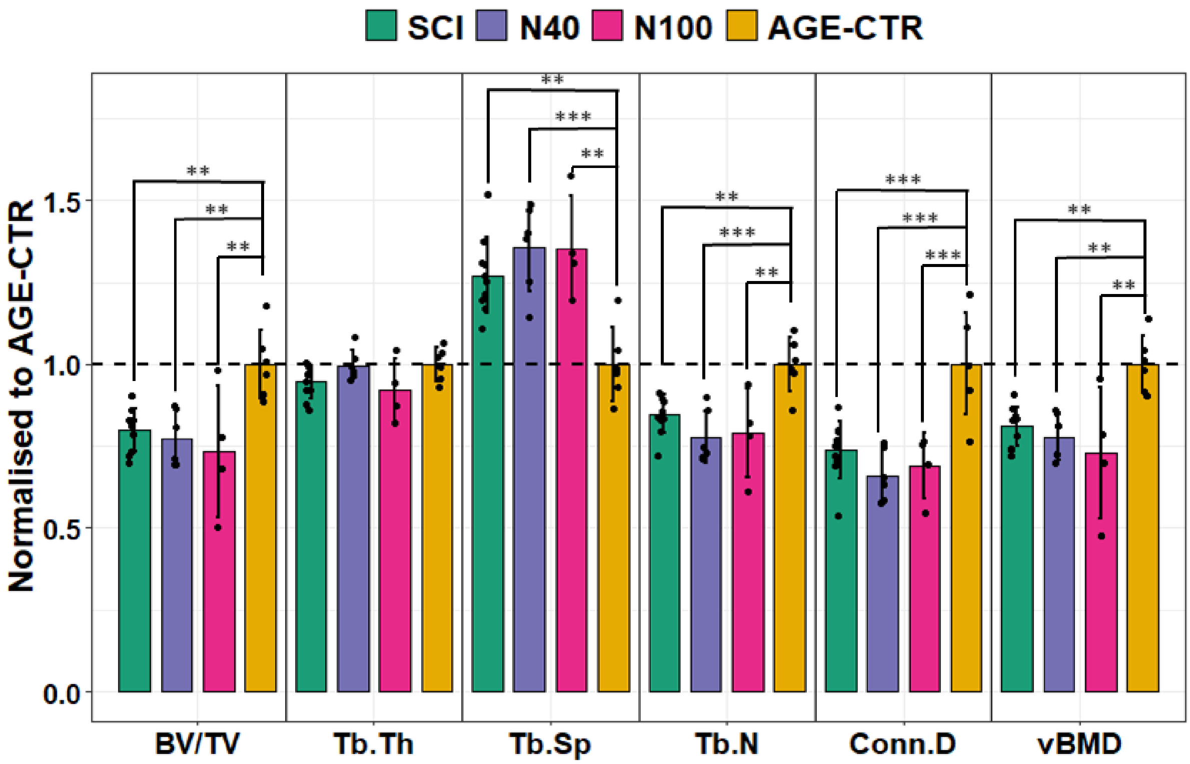


**Figure 8C.** Mean morphometric outcome measures for the proximal tibial epiphyseal secondary spongiosa VOI in the right hindlimbs of SCI, N40, N100 and AGE-CTR rat groups. Data shown as mean ± SD with each parameter normalised to that of AGE-CTR. ** and *** indicate < 0.01 and p < 0.001, respectively.

*ii) Distal Femoral metaphyseal trabecular bone*

- *Contralateral comparison of N40 group*

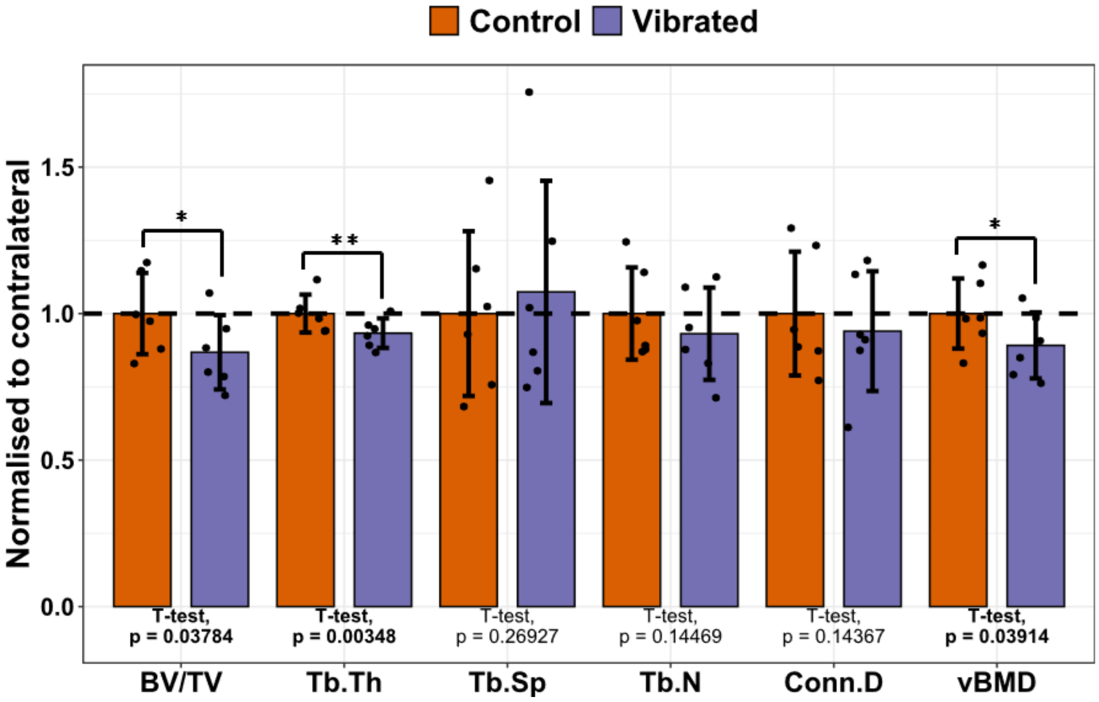


**Figure 8D**. Mean morphometric outcome measures of the nanovibrated (right) and contralateral control (left) distal femur metaphyseal secondary spongiosa VOI for 40 nm amplitude vibrated (N40) rats. Data shown as mean ± SD with each parameter normalised to that of contralateral control. * and ** indicate $p < 0.05$ and $p < 0.01$, respectively.

- *Contralateral comparison of N100 group*

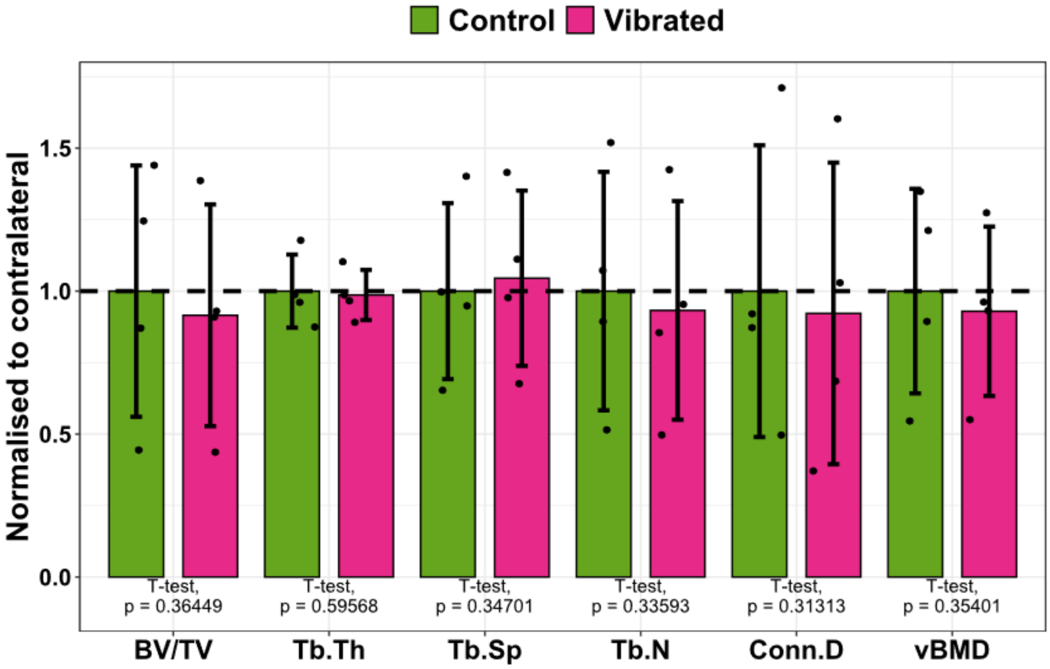


**Figure 8E.** Mean morphometric outcome measures of the nanovibrated (right) and contralateral control (left) distal femur metaphyseal secondary spongiosa VOI for 100 nm amplitude vibrated (N100) rats. Data shown as mean ± SD with each parameter normalised to that of contralateral control.

- *Comparison of nanovibrated/right hindlimb between all groups*

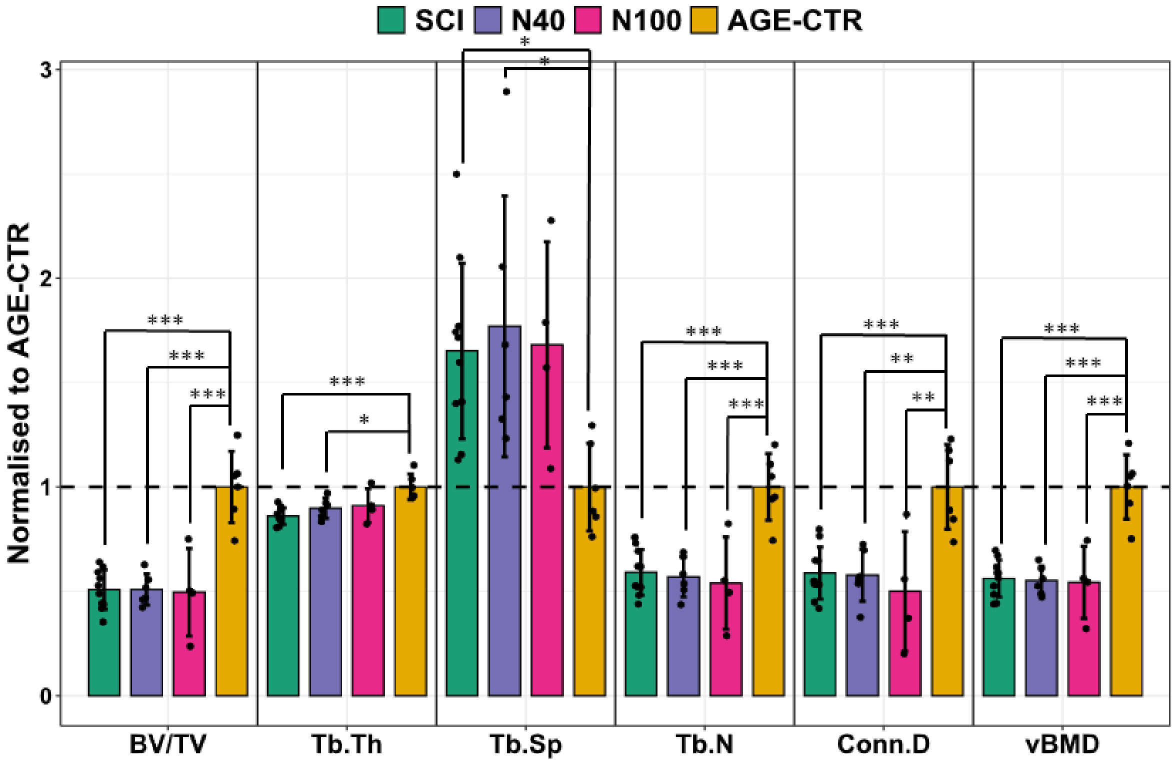


**Figure 8F.** Mean morphometric outcome measures for the distal femur metaphyseal secondary spongiosa VOI in the right hindlimbs of SCI, N40, N100 and AGE-CTR rat groups. Data shown as mean ± SD with each parameter normalised to that of AGE-CTR. *, ** and *** indicate $p < 0.05$, $p < 0.01$ and $p < 0.001$, respectively.

*iii) Distal Femoral epiphyseal trabecular bone*

- *Contralateral comparison of N40 group*

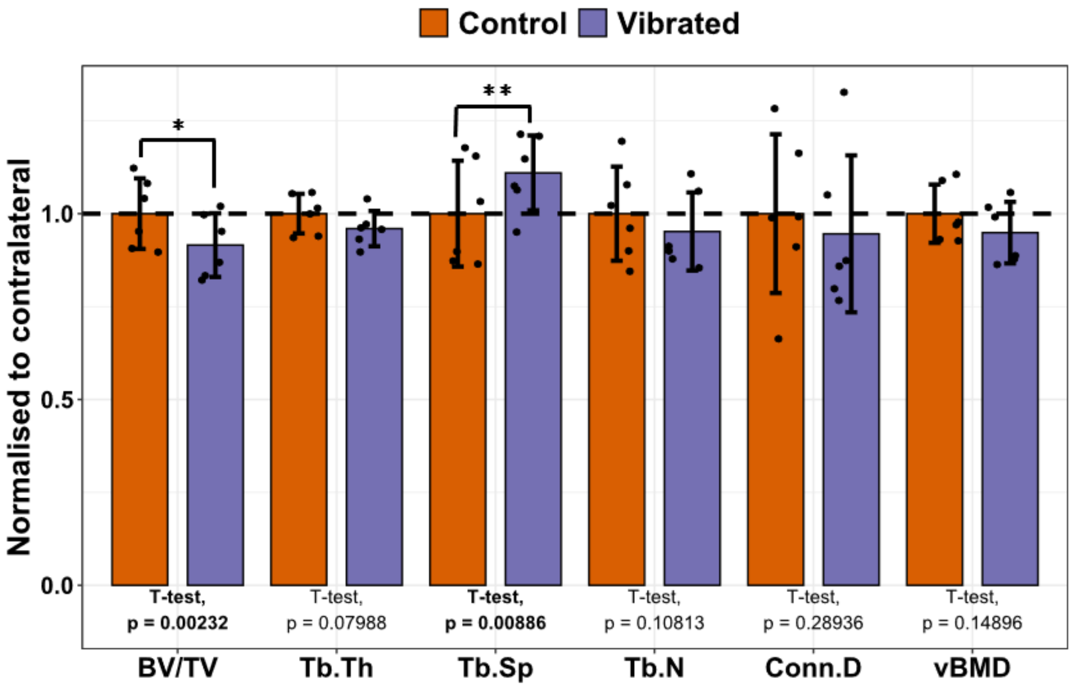


**Figure 8G.** Mean morphometric outcome measures of the nanovibrated (right) and contralateral control (left) distal femur epiphyseal trabecular VOI for 40 nm amplitude vibrated (N40) rats. Data shown as mean ± SD with each parameter normalised to that of contralateral control.

- *Contralateral comparison of N100 group*

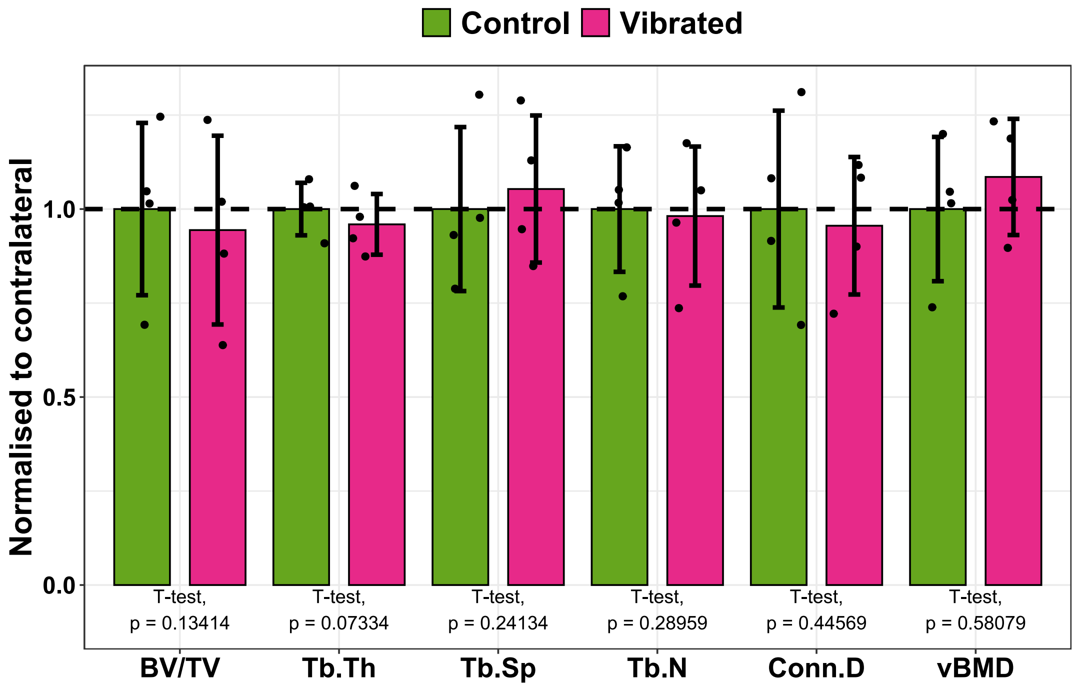


**Figure 8H.** Mean morphometric outcome measures of the nanovibrated (right) and contralateral control (left) distal femur epiphyseal trabecular VOI for 100 nm amplitude vibrated (N100) rats. Data shown as mean ± SD with each parameter normalised to that of contralateral control.

- *Comparison of nanovibrated/right hindlimb between all groups*

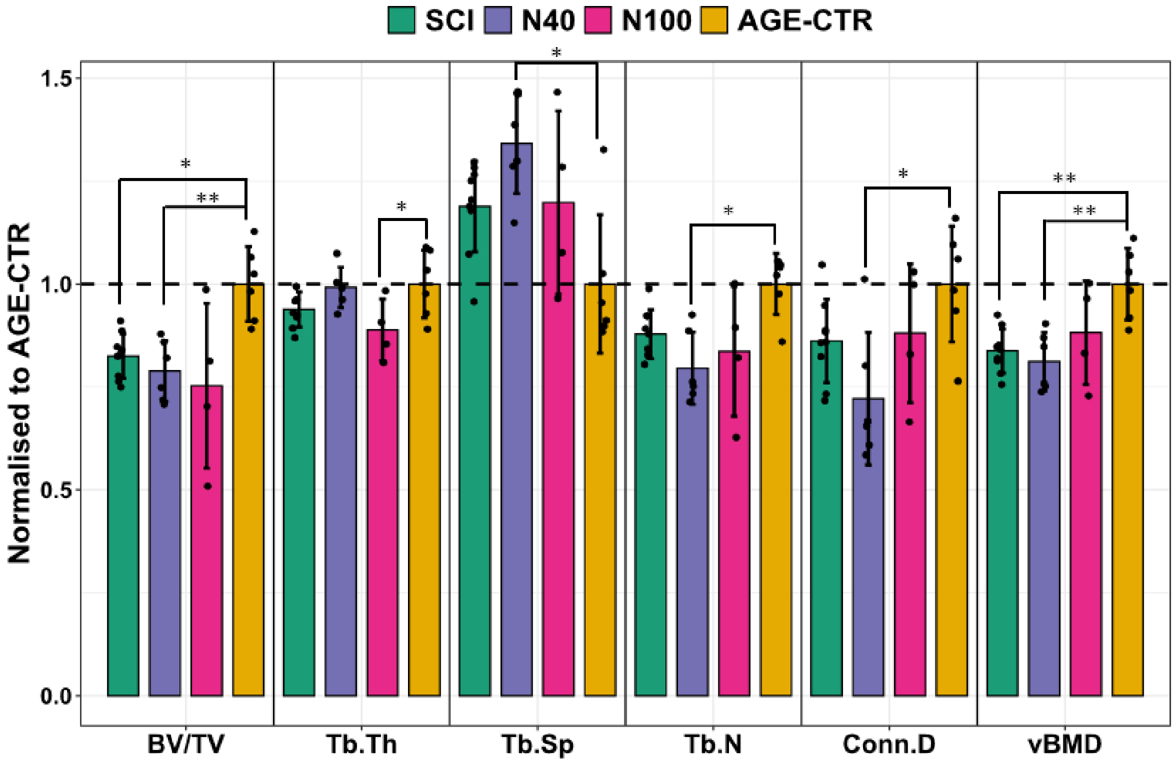


**Figure 8I.** Mean morphometric outcome measures for the distal femur epiphyseal trabecular VOI in the right hindlimbs of SCI, N40, N100 and AGE-CTR rat groups. Data shown as mean ± SD with each parameter normalised to that of AGE-CTR. * and ** indicate $p < 0.05$ and $p < 0.01$, respectively.

**Supplementary Information 9:** μCT analysis of tibial metaphyseal mid-diaphyseal cortical bone morphometry

The parameters assessed were; cortical area (Ct.Ar), total volume enclosed by the periosteum (Tt.V), cortical bone volume (Ct.V) marrow volume (Ma.V), cortical volume fraction (Ct.V/Tt.V), cortical bone surface to volume ratio (BS/BV), cortical bone thickness (Ct.Th), second polar moment of area (J), eccentricity (Ecc) and tissue mineral density (TMD).

- *Contralateral comparison of N40 group*

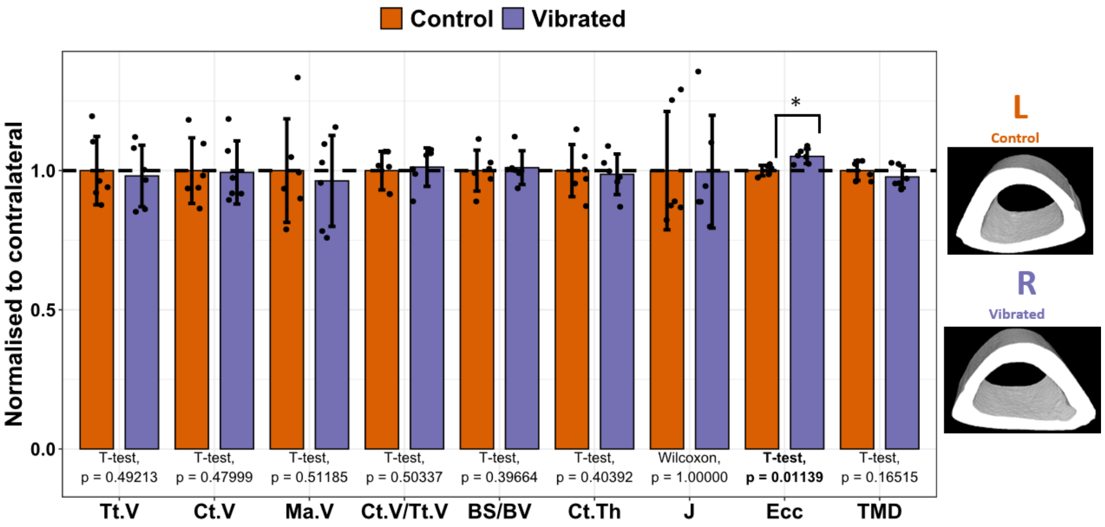


**Figure 9A.** Representative μCT-based images of the nanovibrated (right) and contralateral control (left) tibial mid-diaphyseal cortical bone VOI with mean morphometric outcome measures for 40 nm amplitude vibrated (N40) rats. Data shown as mean ± SD with each parameter normalised to that of contralateral control. * indicates $p < 0.05$.

- *Contralateral comparison of N100 group*

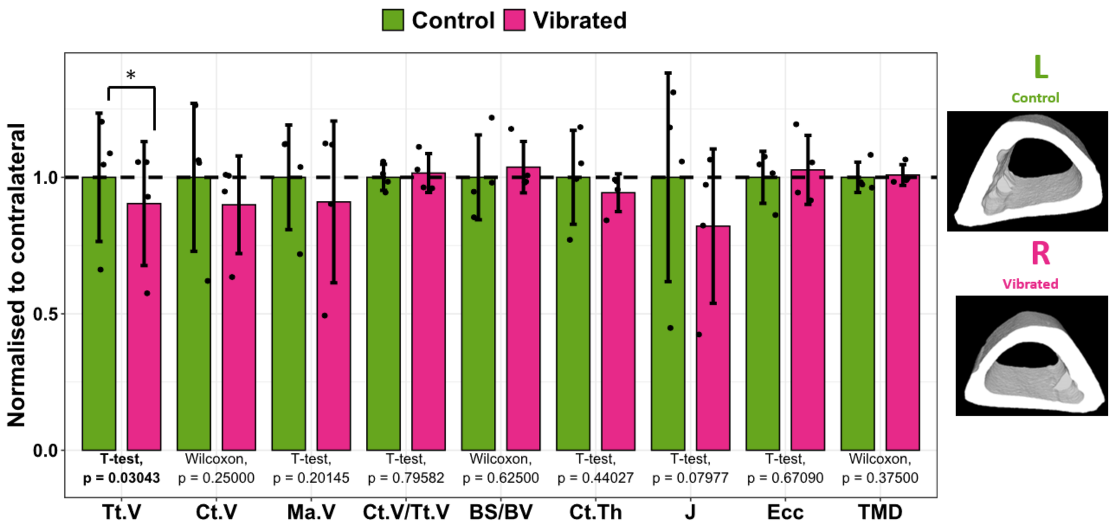


**Figure 9B.** Representative µCT-based images of the nanovibrated (right) and contralateral control (left) tibial mid-diaphyseal cortical bone VOI with mean morphometric outcome measures for 100 nm amplitude vibrated (N100) rats. Data shown as mean ± SD with each parameter normalised to that of contralateral control. * indicates $p < 0.05$.

- *Comparison of nanovibrated/right hindlimb between all groups*

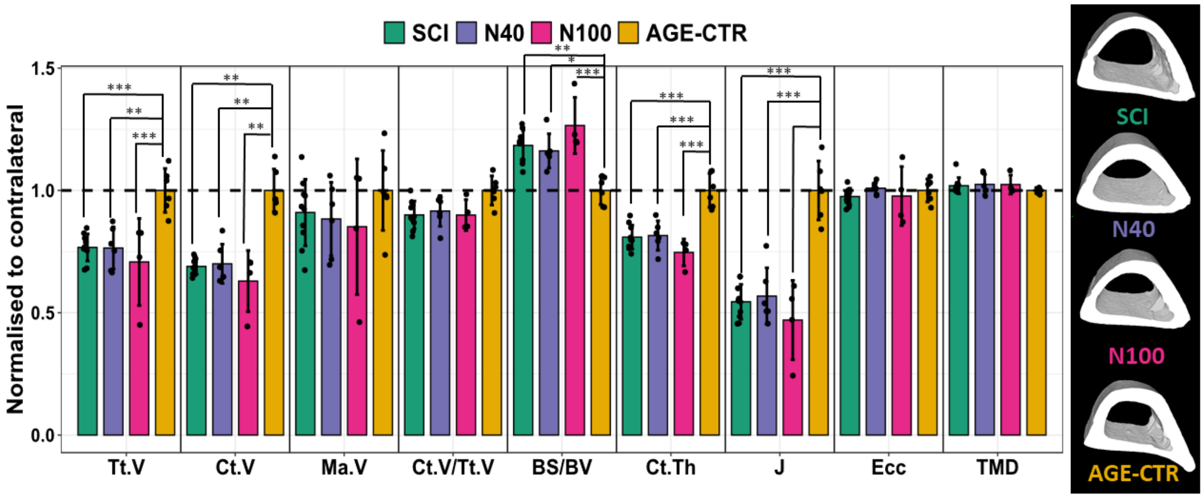


**Figure 9C.** Representative µCT-based images of the tibial mid-diaphyseal cortical bone VOI with mean morphometric outcome measures for SCI, N40, N100 and AGE-CTR rat groups. Data shown as mean ± SD with each parameter normalised to that of AGE-CTR. *, ** and *** indicate $p < 0.05$, $p < 0.01$ and $p < 0.001$, respectively.

**Supplementary Information 10: Three-point bend-determined whole-bone and material-level mechanical properties of tibial mid-diaphyseal cortical bone**

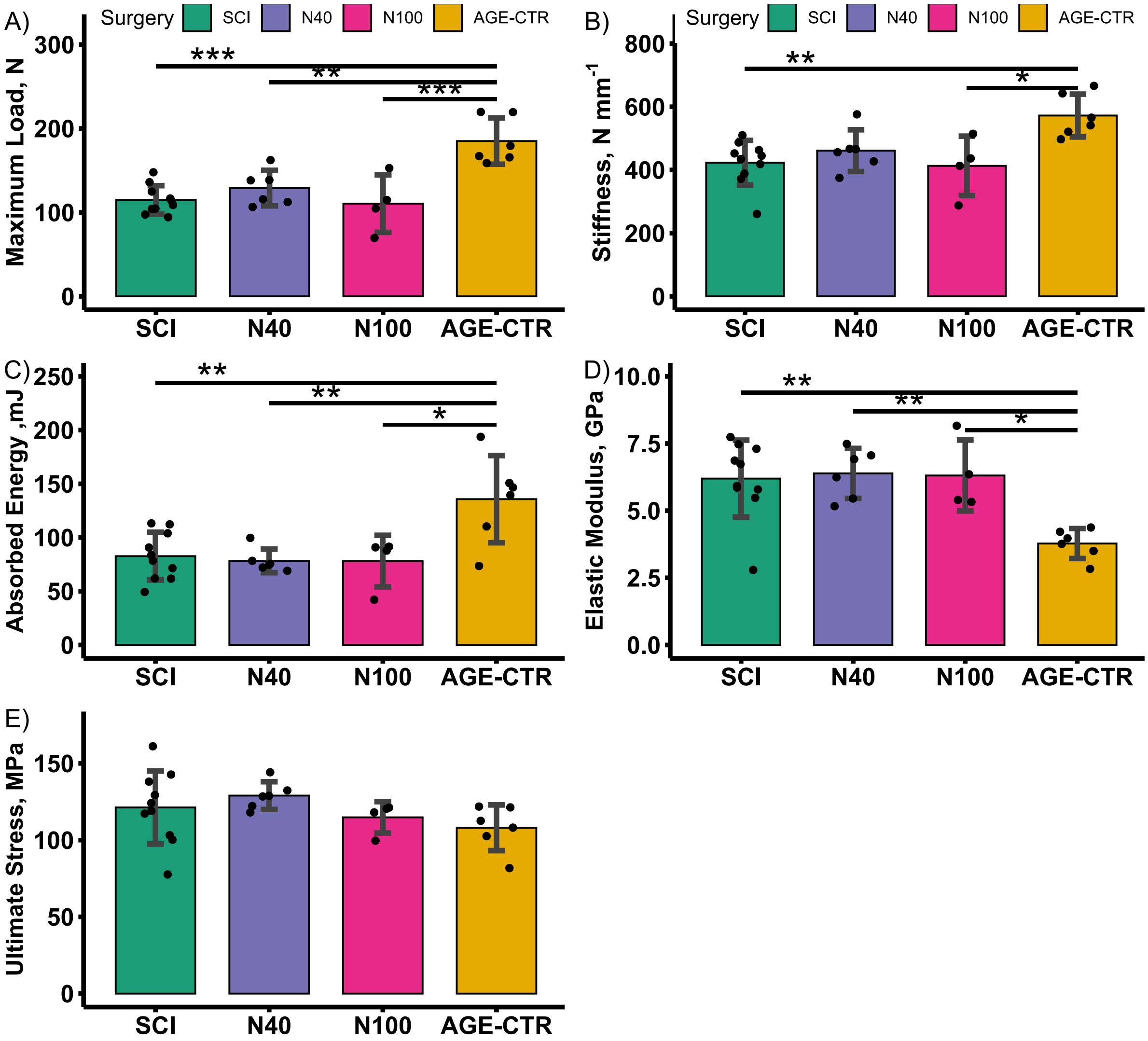


**Figure 10.** Three-point bend-determined whole-bone and material-level mechanical properties for Time-0 and 1-, 2- and 4-weeks post-surgery SCI and SHAM groups. Data shown as mean ± SD. *, ** and *** indicate p < 0.05, p < 0.01 and p < 0.001, respectively.

**Supplementary Information 11: Location of trabecular bone volume of interest (VOI) for degree of anisotropy analysis**

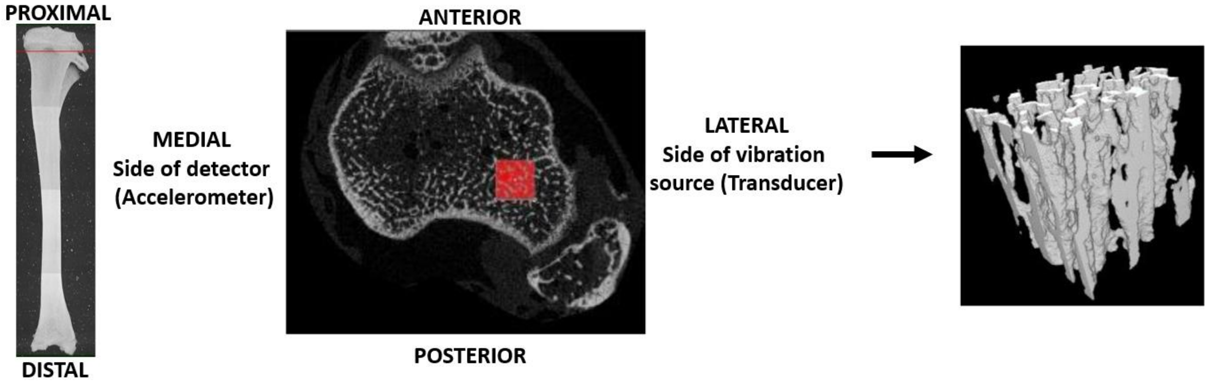


**Figure 11:** Location of top slice of trabecular bone cube subVOI used for degree of anisotropy analysis (in red). Cubic subVOI extends distally for 1.2mm.
